# Phosphorylated ubiquitin is a secondary messenger and an epigenetic mark mediating mitochondria to nucleus signaling

**DOI:** 10.64898/2026.04.24.719390

**Authors:** Thomas J. Mercer, Bence Daniel, Craig Fredrickson, Daniel Le, Serena Lee, Vineet Vinay Kulkarni, Xu Hou, Fabienne C. Fiesel, Hai Ngu, Min Jung, Brent J Ryan, Rachel Heon-Roberts, Amelia J Smith, Vasumathi Kameswaran, Tommy K Cheung, Denise Gastaldo, Dennis W. Dickson, Wolfdieter Springer, Claire Jeong, Oded Foreman, Christopher M. Rose, Baris Bingol

## Abstract

Parkinson’s disease (PD) is commonly associated with dysfunctional mitochondrial homeostasis. PINK1, a S/T kinase mutated in early-onset PD, generates phosphoserine 65 ubiquitin (pS65Ub) on damaged mitochondria facilitating their removal. Here, we show that pS65Ub translocates into the nucleus after generation at damaged mitochondria and is directly attached to substrates by resident E3 ligases. Histone H2A is a major substrate and is modified at lysine 119 (H2AK119) by the polycomb silencer, E3 ligase RING1B. At nucleosomes, pS65Ub simultaneously suppresses RING1B and potentiates H2A deubiquitinases USP16 and USP21. Epigenetic profiling and RNA sequencing reveal that pS65Ub is enriched at the promoters of poorly expressed yet dynamically regulated genes and is associated with H2AK119ub depletion. Functionally, we show that pS65Ub enrichment drives polycomb target gene expression, which accelerates the maturation of dopaminergic neurons. Importantly, post-mortem PD brains exhibit elevated nuclear pS65Ub, potentially linking nuclear pS65Ub accumulation with disease pathogenesis. Together, these data indicate that pS65Ub generated at damaged mitochondria regulates fundamental cellular processes at distant sites.

## Introduction

The mitochondrial damage sensors PINK1 and Parkin are among the most commonly mutated genes in early-onset Parkinson’s disease (PD)^1,2^. In basal conditions, serine/threonine kinase PINK1 is constitutively imported into mitochondria, where it is N-terminally cleaved by resident proteases^3–5^. Cleaved PINK1 then retrotranslocates back into the cytoplasm and is rapidly degraded^6^. Loss of mitochondrial membrane potential prevents import^7^, stabilizing and activating full length PINK1 at the mitochondrial surface via direct interaction with both the translocase of outer membrane (TOM) complex and the outer mitochondrial membrane (OMM)^3,8–11^. Here, PINK1 phosphorylates ubiquitin at serine 65 (pS65Ub), recruiting and activating ubiquitin E3 ligase Parkin which polyubiquitinates OMM proteins. This creates more PINK1 substrates in a feedforward loop^12,13^, which is dampened by a host of deubiquitinases (DUBs), in particular USP30^14–18^. Alongside E3 ligase activation, pS65Ub inhibits mitochondrial DUBs^15,19–21^ and promotes autophagy receptor binding^22^, which combined condemn damaged mitochondria to degradation in lysosomes.

Inefficient mitochondrial quality control is implicated in the earliest stages of PD^1,23^, however the disease is also characterized by a cascade of dysfunctions in proteostasis, inflammation and both transcriptional and epigenetic regulation^1,24–31^. Understanding how these are connected is crucial to understand disease etiology. Advancements in epigenetic technologies have led to the identification of aberrant DNA methylation and histone post-translational modification (two major modes of epigenetic reprogramming) in both PD tissues and experimental models^27–30,32,33^. For example, multiple disease-relevant genes are aberrantly methylated in the PD brain^27,29,32,34^, linked in part to elevated levels of DNA demethylase TET2 which potentiates inflammation-induced dopaminergic neuron death^29^. Additionally, misregulation of histone post translational modification has been identified, both globally and at PD risk genes^33,35^.

The polycomb system is a group of epigenetic repressors that control developmental signaling, including in the brain^36,37^. It comprises polycomb repressive complex 1 (PRC1) and 2 (PRC2), multiprotein epigenetic modifiers that catalyze histone 2A lysine 119 ubiquitination (H2AK119ub) and histone 3 lysine 27 trimethylation (H3K27me3) respectively^38,39^. H2AK119ub and H3K27me3 typically converge with unmethylated CG dinucleotides at gene promoters, forming Polycomb domains^38–42^. H2AK119ub is deposited first and is crucial for polycomb-dependent gene silencing^43–46^. The repression mechanism is incompletely understood, yet may involve H2AK119ub stabilizing both PRC1 and PRC2 at chromatin ^39,45–47^, inhibiting RNA polymerase II^48,49^ and/or promoting chromatin compaction^50^. While polycomb misregulation is implicated in neurodegeneration^51–56^, its relevance in PD is unclear.

Here, we show that, after generation by PINK1 at mitochondria, pS65Ub accumulates in the nucleus where it is attached to chromatin by PRC1, forming a novel epigenetic mark, H2AK119pS65Ub, which is enriched at developmental gene promoters. Mechanistically, pS65Ub potentiates removal of ubiquitin from H2AK119 by simultaneously reducing ligation and promoting deubiquitination. This alleviates transcriptional repression at polycomb target genes and drives functional gene programs. Notably, nuclear pS65Ub is elevated in PD patient brains. Our findings suggest a role for pS65Ub in mitochondria to nucleus signaling, involving a novel secondary messenger-like mechanism and a new mode of substrate pS65 ubiquitination.

## Results

### Phosphoserine 65 Ubiquitin Accumulates in the Nucleus

We aimed to characterize new functions of pS65Ub. First, we examined its subcellular localization in fibroblasts treated with mitochondrial depolarizers commonly used to model dysfunction seen in neurodegenerative diseases^13,57,58^. Despite the well-reported role of OMM-localized pS65Ub, the ionophores carbonyl cyanide 3-chlorophenyl hydrazone (CCCP) and valinomycin, and mitochondrial electron transport chain inhibitors oligomycin/antimycin A (OA) enriched pS65Ub throughout the cell, including in the nucleus (**Extended Data Figure 1A**). Nuclear pS65Ub enrichment increased in a time dependent manner relative to the cytoplasmic pool (**Extended Data Figure 1A**) and was detected using multiple pS65Ub-specific antibodies (**Extended Data Figure 1B**). As we observed similar nuclear pS65Ub accumulation in both iPSC-derived NGN2-cortical (**Figure 1A**) and dopaminergic (**Extended Data Figure 1C**) neurons, we focused on this pool for further examination.

**Figure 1.**
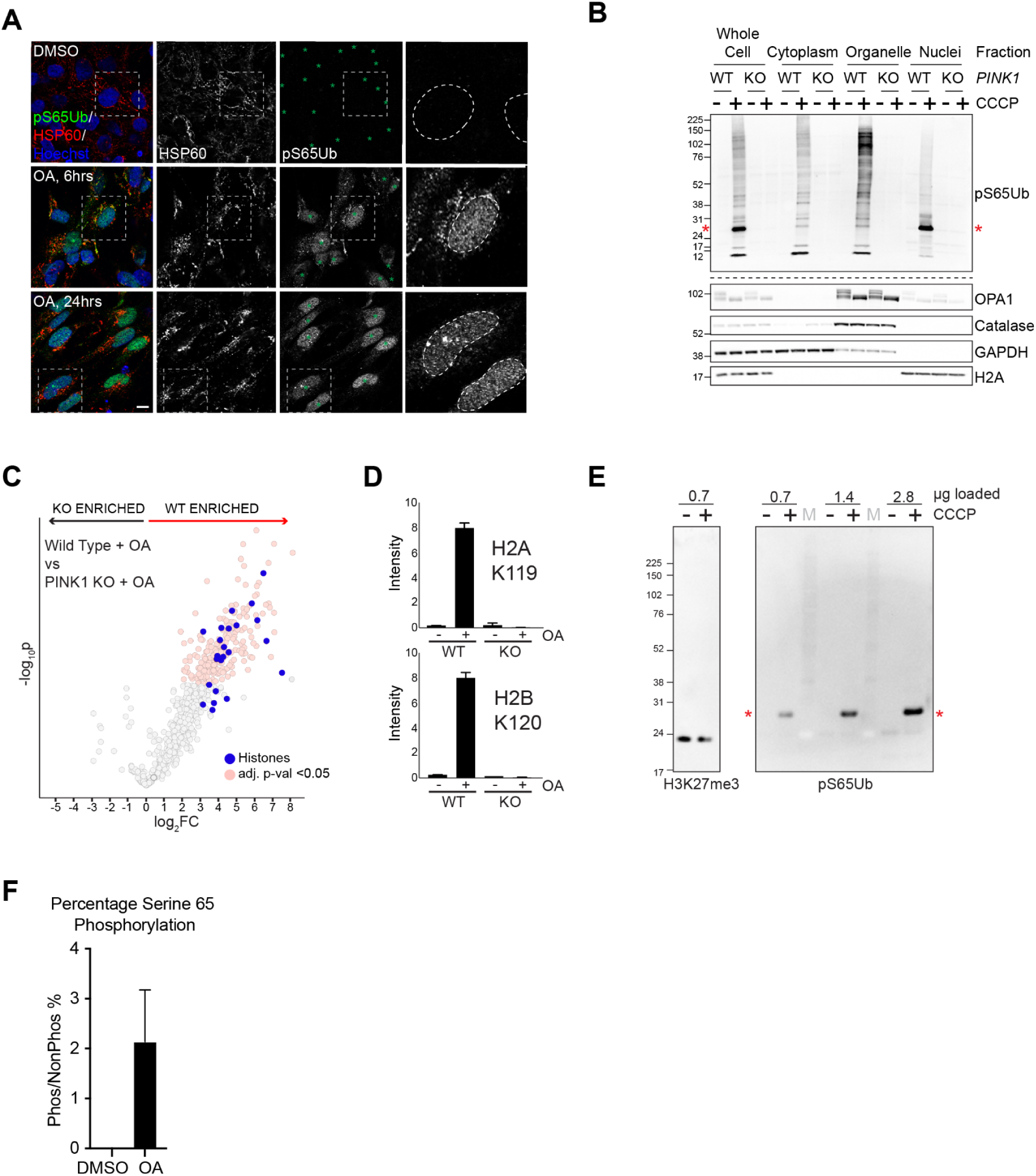
Characterizing the nuclear pS65Ubome. **A** - NGN2-induced iPSC-derived neurons were treated with Oligomycin 1µM + Antimycin 1µM (OA,1µM) for 0, 6 or 24 hrs before visualization of pS65Ub (green), HSP60 (red) and DNA (blue) by immunofluorescence. Dashed grey boxes indicate areas magnified in inset from pS65Ub channel, and nuclei are indicated with green asterisks (dashed circle in inset). Scale bar 10µm. **B** - Wild type (WT) and PINK1 knockout (KO) HEK293T cells were treated with CCCP (20µM, 4 hrs) as indicated. Subcellular fractionation followed by immunoblotting reveals the subcellular distribution of pS65Ub. Red asterisks mark the pS65Ub-histone band. **C** - pS65Ub was immunoprecipitated from nuclear lysates generated from wild type or PINK1 KO HEK293T cells after treatment with OA (1µM, 18 hrs) or DMSO, then samples were subjected to tandem mass tag (TMT)-based quantitative proteomics. Volcano plot showing relative protein enrichment in OA-treated WT vs PINK1 KO cells. Statistically significant proteins are annotated in pink (adjusted p value <0.05) and histones in blue. **D** - Intensity scores from each experimental condition for ubiquitinated sites (identified by searching for KGG-modified peptides in the pS65Ub-enriched samples) mapping to H2AK119 and H2BK120 (mean +/- SEM, three independent repeats). **E** - Histones were purified from CCCP-treated HeLa cells (20µM, 4 hrs) then analysed by Western blot. H3K27me3 was probed to confirm that equal sample amounts were loaded and a red asterisk marks the pS65Ub-histone band. Protein mass marker lanes are marked with the letter M. **F** - The proportion of serine 65-phosphorylated ubiquitin in purified histone fractions from OA- or DMSO-treated HeLa was examined by AQUA proteomics. Mean +/- SEM from three independent repeats.

### Characterizing the Nuclear pS65Ubome

We turned to subcellular fractionation followed by immunoblotting to characterize substrate pS65 ubiquitination via an orthogonal technique. Nuclear pS65Ub accumulation was PINK1-dependent, with immunoblotting for pS65Ub revealing smear in the nuclear fractions indicating its attachment to a range of nuclear substrates (**Figure 1B; Extended Data Figure 1D**). To identify these proteins, we treated wild type (WT) and PINK1 knockout (KO) HEK293T cells with OA, generated nuclear lysates then isolated pS65Ub-proteins via affinity purification before quantitative proteomics. We detected 879 putative nuclear pS65Ub substrates (**Extended Data Table 1**), with gene set enrichment analysis indicating roles in protein translation, response to amino acid starvation and nervous system development, among other roles (**Extended Data Table 2**).

Interestingly, multiple histone proteins were identified (**Figure 1C**; **Extended Data Table 1**), and by searching for ubiquitin conjugation sites (without direct KGG purification), H2AK119 and H2BK120 were identified as putative acceptor residues (**Figure 1D**). We hypothesised that the ∼27kDa pS65Ub band detected in CCCP-treated HEK293T and HeLa cells (**Figure 1B; Extended Data Figure 1D**, red asterisks), which was nucleus-derived and the major single band in the whole-cell lysate, was a mono-pS65 ubiquitinated histone(s). To confirm this, we purified histones from HeLa cells, and showed we were able to isolate a pS65Ub-positive band at the same molecular weight after mitochondrial damage only, referred to herein as the pS65Ub-histone band (**Figure 1E**, red asterisks).

pS65Ub-histones were detected with two independent pS65Ub antibodies (**Extended Data Figure 1E**), and in a diverse range of cell lines including human neurons (**Extended Data Figure 1F, G**). They accumulated in HeLa cells treated with the Parkinson’s disease-associated pesticides Rotenone and Paraquat, and their generation was prevented by cotreatment with PINK1 inhibitor PRT062607^59^ (**Extended Data Figure 1H**). Furthermore, we observed histone pS65 ubiquitination after treatment with the HSP90 inhibitor GTPP, which promotes the mitochondrial unfolded protein response^60,61^, activating PINK1 largely independently of mitochondrial depolarization^61^ (**Extended Data Figure 1I**).

To further characterise the nuclear pS65Ub pool, we used two main toxins: CCCP (to rapidly and maximally depolarize mitochondria) and OA (to gradually induce mitochondrial dysfunction via specific inhibition of oxidative phosphorylation, mimicking chronic disease states more closely), and three main cell types: HeLa and HEK293T cells (due to their high pS65Ub levels, **Extended Data Figure 1F**) and iPSC-derived neurons (due to their physiological relevance). To estimate the stoichiometry of histone pS65Ubiquitination in one of these model systems, we treated HeLa cells with OA then used Absolute QUAntification (AQUA) proteomics to measure the ratio of phosphorylated vs non phosphorylated ubiquitin in purified histone samples. These data suggest that ∼2% of histone-attached ubiquitin is phosphorylated at serine 65 in these conditions (**Figure 1F**).

Together, these data indicate that pS65Ub is attached to a range of nuclear proteins as a generalised response to PINK1 activation. This includes histones, which are the major mono-pS65Ub acceptors in the cell.

### pS65Ub Localizes to Developmental Gene Promoters and is Enriched at Repressed Chromatin

The attachment of pS65Ub to histones led us to investigate its genomic localization upon mitochondrial damage with CUT&RUN (Cleavage Under Targets and Release Using Nuclease)^62^. In DMSO-treated HeLa cells the pS65Ub CUT&RUN peaks were distributed widely across the genome, which we considered as background signal due to the lack of mitochondrially-derived pS65Ub. Strikingly, OA treatment induced promoter-localized peaks (**Figure 2A**, **B; Extended Data Figure 2A**), which were absent with a nonspecific IgG (**Figure 2B**). We identified 4227 genes displaying significant pS65Ub enrichment at promoters in OA-treated cells. The pS65Ub-positive genes (**Extended Data Table 3**) include top hit *BCL11A*, which is implicated in brain development and is specifically expressed in a subset of dopaminergic neurons vulnerable to neurodegeneration in PD^63^ (**Figure 2B**). Gene set enrichment analysis (GSEA) indicated a role of pS65Ub-positive genes in developmental programs (**Figure 2C**, **Extended Data Table 4**). Notably, the pS65Ub gene set was mostly devoid of nuclear-encoded mitochondrial genes^64^ (other than top hit, *EFHD1*), however transcriptional regulators of mitochondrial biogenesis were present including peroxisome proliferator-activated receptor isoforms and coregulators^65^ (**Extended Data Table 3**).

**Figure 2.**
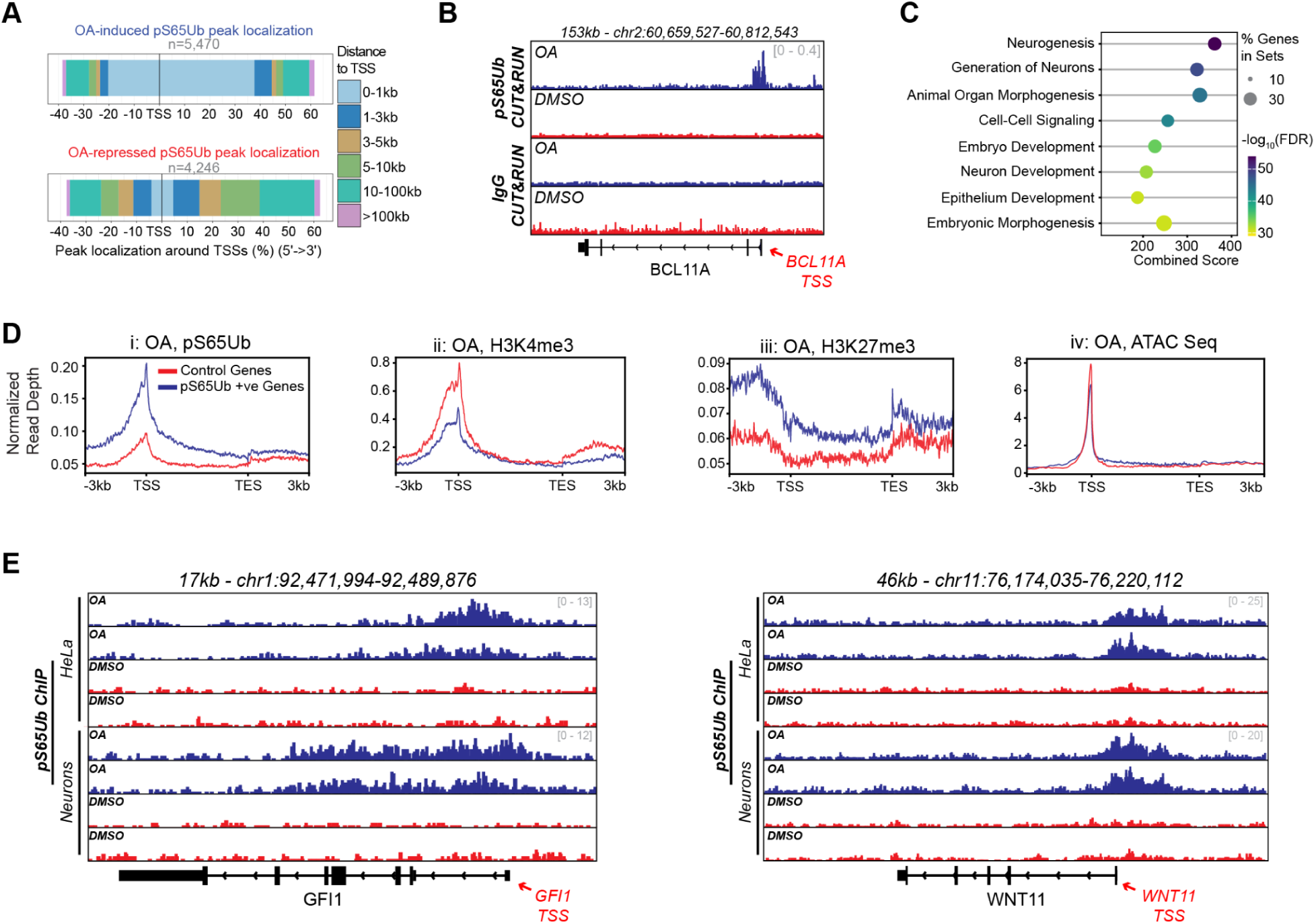
Polycomb target gene promoters are pS65Ub-enriched. **A** - Quantification of pS65Ub CUT&RUN signal relative to the nearest transcription start site (TSS) reveals OA-induced enrichment at the promoter region. **B** - HeLa were treated with OA (1µM, 24 hrs) or DMSO before CUT&RUN analysis with anti-pS65Ub or negative control IgG antibodies. Normalized CUT&RUN peaks from the genomic region spanning BCL11A are shown, with TSS annotated (scale = read coverage). **C** - Bubble plot showing results of gene set enrichment analysis on pS65Ub-positive genes. Bubble color indicates statistical significance of identification and size corresponds to the proportion of matching genes per term. **D** - HeLa cells were treated with OA as before then analysed by CUT&RUN and ATAC Seq. Averaged pS65Ub (i), H3K4me3 (ii) and H3K27me3 (iii) CUT&RUN signal, or ATAC Seq signal (iv) from pS65Ub-positive (blue, n=948) and control gene sets (red, n=1000) genes are plotted separately. Metagene profiles are aligned with gene transcription start (TSS) and end (TES) sites. Experiments i, ii/iii and iv were performed independently. **G** - Normalized pS65Ub ChIP signal from HeLa cells and neurons treated with DMSO or OA as before.Representative genes GFI1 and WNT11 are displayed

We next assessed the distributions of active (H3K4me3) and repressive (H3K27me3) histone marks, and chromatin accessibility, at genes with the strongest pS65Ub promoter signal (mean relative enrichment > 0.5, **Extended Data Table 3**). Compared to 1000 randomly selected genes, pS65Ub-positive genes showed relative H3K4me3 depletion, H3K27me3 enrichment, and lower mean promoter accessibility, all of which are indicative of a repressive epigenetic milieu (**Figure 2D; Extended Data Figure 2B**). We then compared the pS65Ub-positive genes with publicly available chromatin immunoprecipitation sequencing (ChIP Seq) datasets^66^, reasoning that proteins with similar DNA binding profiles (termed coregulators) might function in the same pathways as pS65Ub (**Extended Data Figure 2C**; **Extended Data Table 5**).

Interestingly, the coregulator analysis identified the DNA demethylase TET2 (**Extended Data Figure 2C**, highlighted green), implicated in PD pathogenesis^29,30^ and recently in the regulation of H2AK119ub levels^67^, as a top hit. Transcriptional repressors, including multiple polycomb complex-associated proteins (**Extended Data Figure 2C**, highlighted red) were also enriched suggesting pS65Ub is present at Polycomb domains, corroborating H2AK119 as a pS65Ub acceptor and potentially indicating a role for pS65Ub in epigenetic repression.

### Promoter pS65Ub Enrichment is Observed in Neurons

Next, we compared the genomic distribution of pS65Ub across cell types, focusing on iPSC-derived NGN2-cortical neurons and HeLa cells. CUT&RUN did not yield robust data in mature neurons, likely because extensive neurite branching increased cellular damage when generating single-cell suspensions. We therefore turned to ChIP Seq. Cells were treated with OA (1µM, 24 hrs) before pS65Ub ChIP Seq analysis (with pS65Ub-histone accumulation confirmed in **Extended Data Figure 2D**). We recapitulated OA-dependent promoter-biased pS65Ub enrichment in both HeLa cells and neurons (**Figure 2E**; **Extended Data Figure 2E**), with detection using two chromatin profiling techniques and in two cell types increasing confidence pS65Ub promoter localization. Genes displaying OA-dependent pS65Ub promoter enrichment in the neuron and HeLa cell ChIP Seq experiments are listed in **Extended Data Tables 6 and 7**. We noted a highly significant overlap between cell types (p value < 2.2e-16, Fisher’s exact test) (**Extended Data Figure 2F**; **Extended Data Table 6, 7**), and GSEA analysis of genes common between HeLa and neurons yielded multiple developmental terms (**Extended Data Tables 8-10**) in line with the CUT&RUN dataset (**Figure 2C**; **Extended Data Table 4**).

Together, the enrichment of pS65Ub at developmental gene promoters in distinct cell types is consistent with pS65Ub recruitment to Polycomb domains being a robust and generalized response to mitochondrial damage. Moreover, these data implicate H2AK119 as a nuclear pS65Ub acceptor by an orthogonal approach.

### Histone pS65 Ubiquitination is a Rapid, Stress-Responsive Readout of PINK1 Activation

We went on to characterize how histone pS65 ubiquitination occurs. HeLa cells accumulated pS65Ub-modified histones within 15 min of treatment with 20 µM CCCP (**Figure 3A**). Accumulation was detected at sub-threshold CCCP concentrations associated with incomplete mitochondrial depolarization yet maximal histone modification was seen at higher CCCP concentrations, indicating that histone pS65 ubiquitination scales with the extent of mitochondrial depolarization (**Extended Data Figure 3A, B**). Histone pS65 ubiquitination is preceded by PINK1’s mitochondrial recruitment, as preventing PINK1 stabilisation at the OMM via TOM7 knock down^3^ abolished histone pS65 ubiquitination altogether (**Figure 3B; Extended Data Figure 3C**). These data suggest that histone modification is highly dynamic and responsive to stress-dependent recruitment of PINK1 to mitochondria.

**Figure 3.**
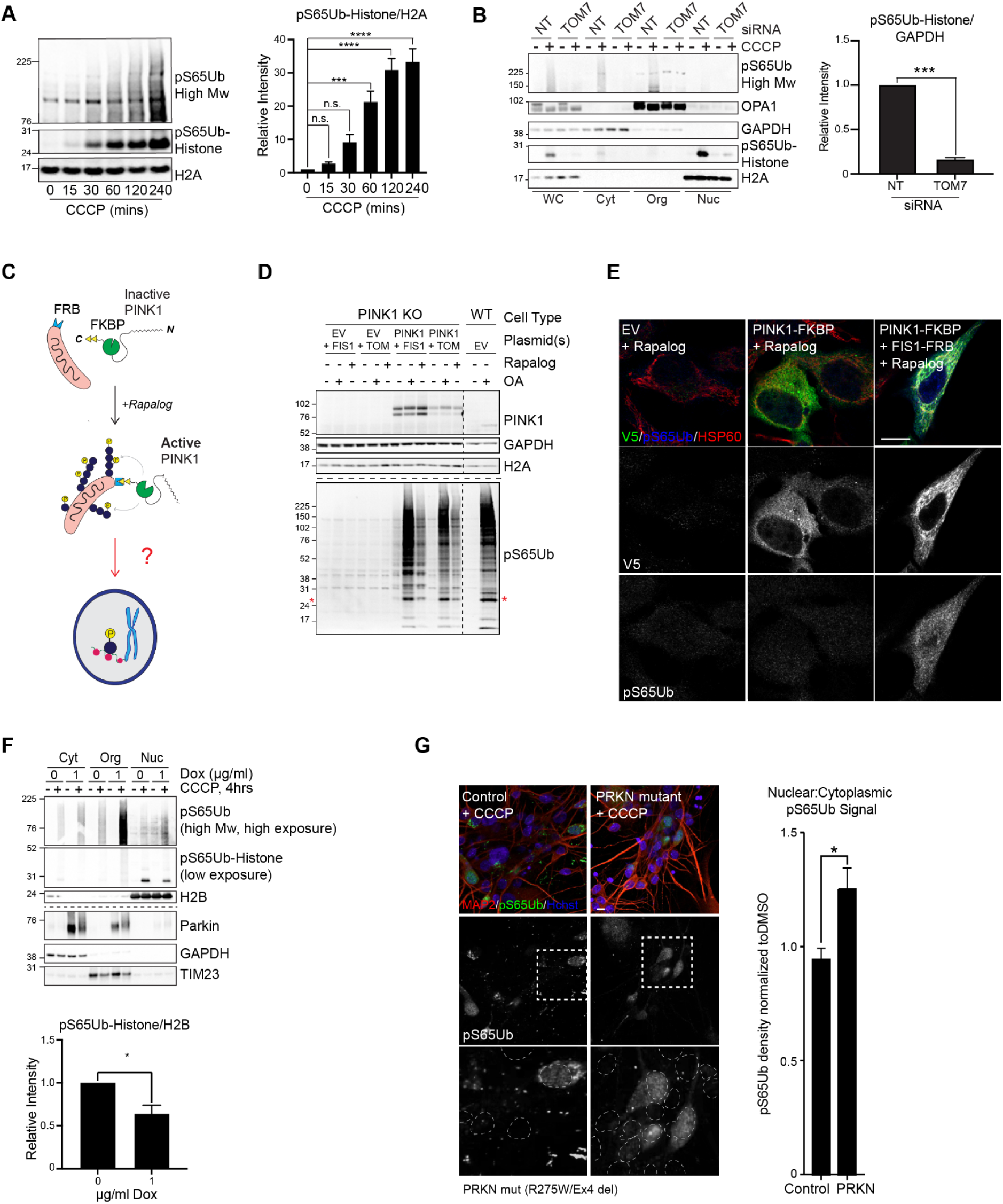
Mitochondrially-derived pS65Ub undergoes nuclear translocation. **A** - HeLa cells were treated with CCCP (20µM) for 0-240 mins before subcellular fractionation and analysis of the nuclear fraction by Western blot. Mean relative intensity of pS65Ub-histone/H2A is plotted, +/- SEM, one-way ANOVA. **B** - Cells transfected with TOM7 or non-targeting (NT) siRNAs were treated with CCCP (20µM, 4 hrs) before subcellular fractionation (WC = whole-cell, Cyt = cytoplasm, Org = membrane-bound organelles, Nuc = nuclei) and Western blot analysis. Mean +/- SEM pS65Ub-histone/GAPDH from the whole-cell fraction was quantified, 1 sample t-test. **C** - Schematic depiction of experiment performed in Figure 3D, E and Extended Data Figure 4D. **D** - HeLa cells were transiently transfected with empty vector (EV), PINK1-FKBP, or PINK1-FKBP and FIS1-FRB then treated with rapalog (500nM, 18 hrs). Cells were fixed and V5 (green), pS65Ub (blue) and HSP60 (red) were visualized by immunofluorescence. **E** - WT or PINK1 KO HeLa were transiently transfected with empty vector (EV), PINK1-FKBP, FIS1-FRB or TOM7-FRB in combinations indicated before 18 hrs treatment with OA (1µM) or rapalog (500nM). The asterisks indicate the position of the pS65Ub-histone band. Dashed lines reveal either where identical samples were analysed on separate gels (horizontal), or where blots were cropped for presentation (vertical). **F** - HEK293T cells expressing a dox-inducible GFP-Parkin cassette were treated with doxycycline for 48 hrs and either CCCP (20µM, 4 hrs) or DMSO before subcellular fractionation and Western blot analysis. pS65Ub/GAPDH levels were quantified from five independent repeats, mean +/- SEM, 1-sample t-test. **G** - iPSC-derived dopaminergic neurons from PD patients with mutations in *PRKN* as well as those from healthy individuals were treated with CCCP (10µM, 6 hrs). Dashed boxes annotate the area magnified in the inset and dashed circles show nuclear borders. Inset zoom in of pS65Ub channel. Quantification shows change in DMSO-normalized nuclear:cytoplasmic pS65Ub density in CCCP-treated neuronal cultures from PD patients with *PRKN* mutations and control individuals. Nuclear and cytoplasmic pS65Ub enrichment in DMSO- and CCCP-treated conditions are plotted in Extended Data Figure 4H. Three independent cell lines per genotype, mean +/- SEM, two sample t test. *p<0.05, ***p<0.001, ****p<0.0001, n.s. not significant, scale bars 10µm.

We next assessed the relationship between histone pS65 ubiquitination and mitophagy in cells exposed to chronic mitochondrial injury. To measure mitophagy rates, we tracked turnover of the mitochondrial inner membrane protein TIM23^68^. As exogenous Parkin expression is required for CCCP-dependent mitophagy in HeLa cells^69^, we performed the experiment in HEK293T cells, which express Parkin endogenously^70–73^. Histone pS65 ubiquitination was detected at 30 mins, peaking at 8 hrs then decreasing at 16 hrs, while TIM23 levels were stable at 30 mins, then monotonically decreased from 4-16 hours, in line with prolonged depolarisation being required for pronounced mitophagy induction^68,70,74^ (**Extended Data Figure 3D**). These data support that histone-pS65Ubiquitination corresponds with PINK1 activation, initially preceding, then occurring in parallel with mitophagic clearance.

### Mitochondrial PINK1 Regulates the Nuclear pS65Ub Pool

PINK1 exists in full length (64kDa) and cleaved (53kDa) proteoforms. Cleaved PINK1 is released from the TOM complex into the cytoplasm where it is constitutively turned over via the ubiquitin-proteasome pathway, preventing kinase accumulation in basal conditions^4–6^. Despite this, cleaved PINK1’s kinase activity and physiological functionality have been reported by multiple groups^75–77^. As exogenously expressed cleaved PINK1 can access the nucleus^78^, we tested whether, after activation at mitochondria, cleaved PINK translocates to the nucleus to modify nucleosomes in situ.

To examine which proteoform drives histone pS65 ubiquitination, we stabilized full length PINK1 via OA treatment and cleaved PINK1 using the proteasome inhibitor MG132, before subcellular fractionation (**Extended Data Figure 4A, B**). Full length PINK1 accumulated in the membrane-bound organelle fraction along with mitochondrial markers, while cleaved PINK1 showed nuclear enrichment. However, pS65Ub-histones only accumulated in the presence of OA, being absent upon MG132 treatment (**Extended Data Figure 4B**, dashed box). Immunofluorescence analysis of cells overexpressing PINK1-HA corroborated these findings, with nuclear pS65Ub only present upon OA treatment and nuclear PINK1 exclusive to MG132 (**Extended Data Figure 4C**). Thus, histone pS65 ubiquitination is only induced by mitochondrial damage (where no nuclear PINK1 is detected), and nuclear cleaved PINK1 is only induced by proteasome inhibition (where no pS65Ub-histones are detected).

To validate full length PINK1 as the pS65Ub-histone-relevant proteoform, we examined the consequence of PINK1 OMM recruitment in the absence of mitochondrial damage signals. Fusion proteins containing FKBP and FRB domains rapidly heterodimerize in the presence of rapamycin analogs (rapalogs)^10^. While rapalog treatment alone was insufficient to activate PINK1-FKBP (**Extended Data Figure 4D**), it induced both nuclear and histone pS65Ub accumulation in cells coexpressing an FRB-tagged OMM localized protein (either FIS1 or TOM7) (**Figure 3C**-**E**). We then aimed to rule out the possibility that a small amount of cleaved PINK1 is liberated from damaged mitochondria to phosphorylate ubiquitin in the nucleus. To control cleaved PINK1’s subcellular localisation, we added C-terminal nuclear export (NES) or import (NLS) signals then assessed nuclear pS65 ubiquitination. While cleaved PINK1 relocalized as expected, neither tag affected the accumulation of pS65Ub in the nucleus or on histones upon CCCP treatment, confirming that cleaved PINK1 nuclear translocation is dispensable for nuclear pS65 ubiquitination (**Extended Data Figure 4E**-**G**).

Collectively, these findings reveal that the activation of PINK1 at mitochondria is sufficient to drive histone pS65 ubiquitination, and that PINK1’s nuclear translocation is neither necessary nor sufficient.

### Parkin Biases pS65 Ubiquitination Toward Mitochondria

Due to its integral role in damage-dependent mitochondrial pS65Ub enrichment, we further dissected how Parkin activity influences histone pS65Ub accumulation. In CCCP-treated HEK293T cells, GFP-Parkin overexpression increased pS65Ub levels in the mitochondria-containing fraction while decreasing histone modification (**Figure 3F**), suggesting that Parkin activity might bias pS65Ub towards mitochondria thus preventing histone modification.

We then assessed the Parkin loss-of-function phenotype in PD patient-derived cells, focusing first on dopaminergic neuronal cultures from PD patients with inactivating mutations in *PINK1* (Q456X/Q456X; Q456X/Q456X; Q456X/Q456X) or Parkin/*PRKN* (c1072tdel/Ex7del; R275W/Ex4 del; R275T/C352R), and using nuclear pS65Ub enrichment as a proxy for histone modification. Nuclear:cytoplasmic pS65Ub signal was significantly increased in the *PRKN* mutant carrier neurons compared to controls, driven by a decrease in the cytoplasmic relative to the nuclear pool (**Figure 3G**; **Extended Data Figure 4H, I**). An increase in nuclear:cytoplasmic pS65Ub was recapitulated in skin fibroblasts bearing *PINK1* (Q456X/Q456X) or *PRKN* (Ex4-7 del/c.203_204 del AG) PD mutations (**Extended Data Figure 4J, K**), where the time-dependent increase in nuclear:cytoplasmic pS65Ub signal observed upon mitochondrial depolarization (**Extended Data Figure 1A**) was enhanced without functional Parkin (**Extended Data Figure 4K**). As expected, PINK1 was required for pS65Ub generation upon mitochondrial damage in both cell types (**Extended Data Figure 4H-K**).

Together, these data suggest that Parkin activity biases pS65 ubiquitination to cytoplasmic targets, likely OMM proteins in accordance with previous reports^13,58,79,80^, and that this may correspond with decreased histone pS65Ubiquitination.

### Histones are Noncanonically pS65 ubiquitinated Inside the Nucleus

As nuclear pS65Ub ultimately derives from damaged mitochondria, we surmised it rapidly traffics to the nucleus where it is attached to histones, potentially akin to a secondary-messenger like system. To test this, we introduced exogenous, monomeric pS65Ub into the cytoplasm of PINK1 KO HeLa cells via electroporation, then compared histone modification over time. Exogenous pS65Ub was rapidly ligated to histones, peaking at 15 mins post-electroporation and tailing off over 2 hrs, with turnover slower for an unphosphorylated control (HA-Ub) (**Figure 4A**). Polyubiquitin levels were unaffected by electroporation suggesting that exogenous ubiquitin did not markedly increase the total pool. We note that the stoichiometry of high Mw (>52kDa) to histone pS65Ub substrates was similar after pS65Ub electroporation and CCCP-treatment, indicating that pS65Ub electroporation phenocopies mitochondrial depolarization with regards to nuclear/histone pS65 ubiquitination (**Figure 4A**).

**Figure 4.**
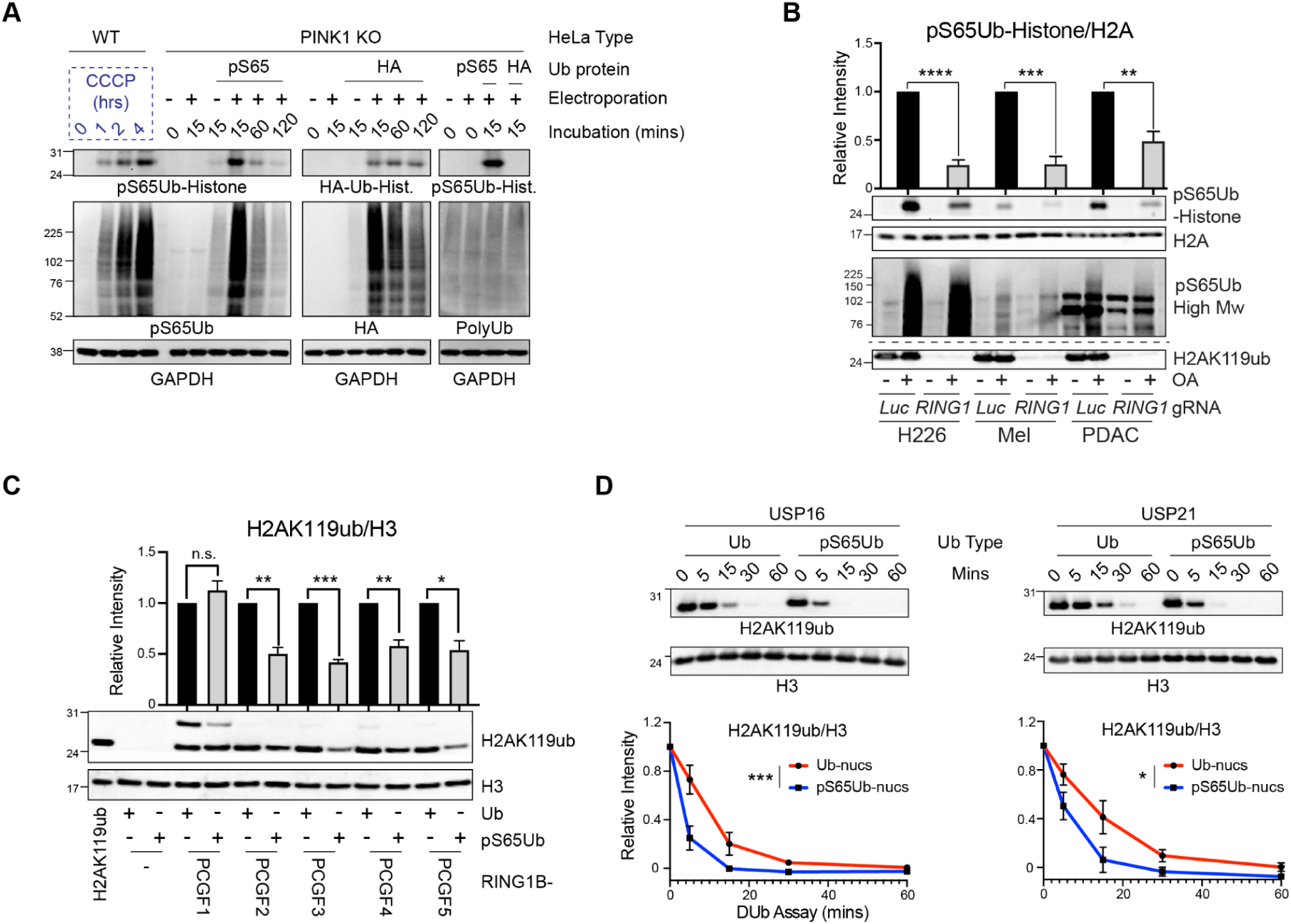
pS65Ub reciprocally inhibits H2AK119 ubiquitination and potentiates deubiquitination. **A** - PINK1 KO HeLa cells were electroporated with monomeric pS65Ub, HA-Ub or in the absence of exogenous ubiquitin, then incubated for 0-120 mins as indicated. In parallel, WT HeLa were treated with CCCP for 0-4 hrs. Whole-cell lysates were analysed by Western blot. The discrete HA-positive ∼27kDa band was annotated as a histone based on similarity to the pS65Ub-histone band. **B** - NCI-H226 (H226), murine melanoma (Mel) or murine pancreatic ductal adenocarcinoma (PDAC) cells were transfected with Cas9 and gRNAs against RING1A and RING1B (*RING1*), or luciferase (*Luc*, negative control) before treatment with OA (1µM, 18 hrs) and analysis by Western blot. Dashed lines indicate where identical samples were analysed on a separate gel. pS65Ub/H2A levels were quantified from 6 (H226) or 5 (Mel/PDAC) independent repeats, with mean +/-SEM and significance values from 1 sample t-tests shown. **C** - RING1B heterodimerised with PCGF1-5 was used to modify mononucleosomes at H2AK119 with either ubiquitin (Ub) or pS65Ub. Nucleosomes pre-ubiquitinated at H2AK119 were used as a positive control. *In vitro* enzymatic efficiency was inferred by measuring relative H2AK119ub accumulation via Western blot. Quantification of H2AK119ub/H3 from four independent repeats is shown. Mean +/-SEM, 1 sample t-test. **D** - H2AK119-ubiquitinated or pS65 ubiquitinated mononucleosomes were incubated with immunopurified deubiquitinases USP16 or USP21 for 0-60 mins as indicated. Mean +/- SEM H2AK119ub/H3 levels from 4 independent experiments are shown below. The effect of ubiquitin type on reaction rate was significantly different by 2 way ANOVA. *p<0.05, **p<0.01, ***p<0.001, ****p<0.0001, n.s. = not significant.

These data imply that two modes of substrate pS65 ubiquitination exist: canonical, where PINK1 phosphorylates pre-conjugated ubiquitin to form pS65Ub^15,20^, and noncanonical, where pS65Ub, generated initially by PINK1 at the OMM, is directly ligated to substrates, likely by a ubiquitin E3 ligase(s). The discovery of cellular evidence for noncanonical pS65 ubiquitination was particularly striking given that pS65Ub itself is not used for chain assembly by Parkin^20,81–83^. So, to further validate noncanonical pS65 ubiquitination, we looked for a pS65Ub-competent E3 ligase, focusing on our strongest candidate substrate, H2AK119.

The main E3 ligase for H2AK119 is PRC1, a multiprotein E3 ligase complex centered on a catalytic core of RING1A or RING1B (collectively RING1) bound to 1 of 6 polycomb group ring finger (PCGF) proteins^38,39^. To test whether PRC1 is necessary for pS65Ub-histone accumulation, we ablated RING1A and RING1B expression in 1 human and 2 murine cancer cell lines, treated them with OA, then assayed pS65 ubiquitination by Western blot. RING1 knockout abolished H2AK119 monoubiquitination and reduced pS65Ub-histone levels by ∼50-70% (**Figure 4B**), consistent with PRC1 being required for the majority of histone pS65 ubiquitination. Next, we reconstituted the reaction *in vitro* using RING1B heterodimerized with PCGF1-5^84^. As the H2AK119ub antibody epitope was unaffected by ubiquitin serine 65 phosphorylation (**Extended Data Figure 5A**), we used H2AK119ub immunoblotting to measure E3 ligase activity. All 5 RING1B-PCGF heterodimers ligated pS65Ub to nucleosomes, however ligase efficiency was reduced compared to unphosphorylated ubiquitin (**Figure 4C**; **Extended Data Figure 5B**).

These results indicate that PRC1/RING1 is capable of noncanonical pS65 ubiquitination, and support H2AK119 as a mono-pS65Ub acceptor in multiple cell types. Since PRC1 acts in the nucleus, they imply that noncanonical pS65 ubiquitination occurs at distal sites after pS65Ub generation at mitochondria by PINK1. Finally, these data suggest that pS65Ub decreases RING1B E3 ligase activity as a suboptimal substrate.

### pS65Ub Counteracts H2AK119 Ubiquitination

We decided to further characterize the consequences of H2AK119 pS65 ubiquitination on nucleosomes. Compared to unmodified ubiquitin, pS65Ub is a suboptimal substrate for most DUBs^15,19,20,85,86^ and stabilization of OMM-localized ubiquitin chains may be one of the mechanisms by which pS65Ub promotes mitophagy^15,16^. We therefore asked whether pS65Ub-histones were resistant or vulnerable to DUB activity, focusing on two H2AK119 DUBs: USP16^87^, which shares gene targets with polycomb group deubiquitinase BAP1^88^, and USP21^89^, which is specifically upregulated among DUBs in PD^90^. We were unable to reconstitute BAP1 activity *in vitro* so excluded it from our analysis.

To compare the impact of ubiquitin S65 phosphorylation on histone deubiquitination, modified nucleosomes were incubated with immunopurified HA-USP16 or HA-USP21 for 0-60 mins, then H2AK119ub levels were tracked to reveal DUB activity. Surprisingly, and in contrast to its effect on RING1B, ubiquitin S65 phosphorylation increased the activity of both USP16 and USP21 on nucleosomes (**Figure 4D**; **Extended Data Figure 5C**). Together, these data suggest that pS65Ub antagonises H2AK119 monoubiquitination by reciprocally reducing PRC1 E3 ligase activity while potentiating DUBs.

### RYBP Recruitment is Unaffected by Ubiquitin Serine 65 Phosphorylation

Finally we looked for potential epigenetic readers by identifying specific pS65Ub-histone interactors^39^. Non-ubiquitinated, ubiquitinated and pS65 ubiquitinated mononucleosomes were immobilized on beads and incubated with HeLa nuclear lysates before identification of coimmunoprecipitating proteins by mass spectrometry.

We were unable to detect any pS65Ub-specific binders, however RYBP, a PRC1 component that binds H2AK119ub to stabilize the complex at chromatin^39,44,47^, was enriched equally by ubiquitinated and phosphoubiquitinated nucleosomes (**Extended Data Figure 5D-F**; **Extended Data Table 11**). These data suggest that pS65Ub does not affect RYBP-dependent PRC1 stabilisation on chromatin.

### Stress-Induced Chromatin-Bound pS65Ub Associates with H2AK119ub Depletion and Gene Activation

H2AK119ub is a repressive epigenetic mark. Therefore, we speculated that pS65Ub-dependent H2AK119ub removal may drive gene expression. To test this hypothesis, we examined H2AK119ub enrichment at OA-induced pS65Ub peak loci in HeLa cells. As expected, pS65Ub and H2AK119ub colocalised (**Figure 5A**), corroborating insights from the nuclear pS65Ubome (**Figure 1C**, **D**), coregulator analysis (**Extended Data Figure 2C**) and RING1 knockout/*in vitro* reconstitution (**Figure 4B**, **C**) experiments. Furthermore, pS65Ub and H2AK119ub were reciprocally regulated, with the OA-induced pS65Ub peaks tending to be depleted of H2AK119ub relative to DMSO (**Figure 5A**). These data suggest that at these loci, while the proportion of pS65Ub is increased, overall H2AK119 modification levels are reduced. In contrast, H2AK119ub was depleted at DMSO-specific pS65Ub peaks, irrespective of treatment condition (**Extended Data Figure 6A**).

**Figure 5.**
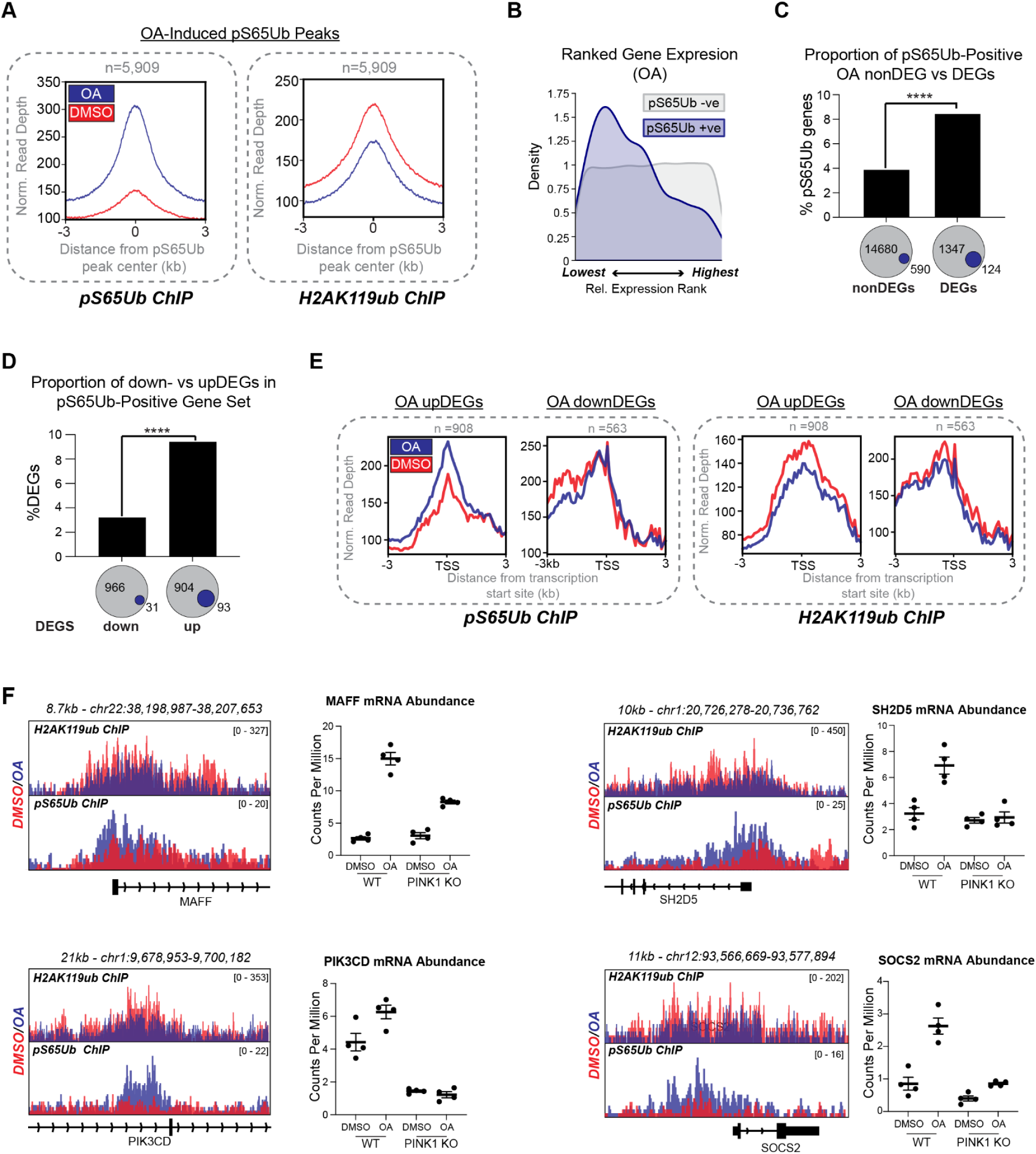
H2AK119ub depletion at pS65Ub-positive promoters correlates with increased gene expression. **A** - pS65Ub ChIP Seq peaks showing induction on OA treatment were subsetted, with averaged pS65Ub and H2AK119ub signal from DMSO (red) and OA (blue) treated HeLa shown. pS65Ub-enrichment correlates with H2AK119ub depletion. **B** - HeLa were treated with OA (1µM, 24 hrs) before RNA Seq analysis. Kernel density estimate plot shows genes ranked by relative expression level, color coded based on presence (blue) or absence (grey) in CUT&RUN geneset. Genes with mean relative enrichment score >0.5 in Extended Data Table 3 were considered pS65Ub-positive. **C** - pS65Ub-positive genes are overrepresented among OA-dependent differentially expressed genes (DEGs). Absolute numbers of pS65Ub-positive (blue) and negative (grey) DEGs and non-DEGs are given below. Fisher’s Exact Test, odds ratio 2.29. **D** - OA-dependent upregulated genes (upDEGs) are significantly enriched relative to downDEGs among pS65Ub-positive genes. Fisher’s Exact Test, odds ratio 3.20. **E** - Metagene profiles showing averaged pS65Ub and H2AK119ub ChIP Seq signal at the transcription start site (TSS) of genes upregulated or downregulated upon OA treatment in WT HeLa (Extended Data Table 12). Both pS65Ub enrichment and H2AK119ub depletion are observed exclusively in the upDEG geneset. **F** - Candidate pS65Ub-dependent genes displaying OA-dependent transcriptional upregulation and both pS65Ub enrichment and H2AK119ub depletion by ChIP Seq (scale units = reads per kilobase per million), as well as reduced transcriptional upregulation upon PINK1 knockout. ** p<0.01, **** p<0.0001.

To test whether pS65Ub-dependent H2AK119ub deubiquitination regulates gene expression, we performed RNA Seq in OA-treated WT HeLa cells (**Extended Data Table 12, Extended Data Figure 6B**). The pS65Ub-positive gene set was generally poorly expressed compared to the remaining non-pS65Ub-enriched genes (**Figure 5B**; **Extended Data Figure 6C**) (in line with the epigenetic indicators described in **Figure 2D**). However, OA-dependent differentially expressed genes (DEGs) were twice as likely to be pS65Ub-positive compared with nonDEGs. (**Figure 5C, Extended Data Figure 6D**), and pS65Ub-positive DEGs showed a strong bias towards upregulation (**Figure 5D**; **Extended Data Figure 6B, E**). Strikingly, OA-dependent pS65Ub enrichment and H2AK119ub depletion was observed exclusively in the upregulated gene set (**Figure 5E**; **Extended Data Table 12**), suggesting that pS65Ub may activate gene expression by relieving H2AK119ub-dependent gene repression.

To test this hypothesis further, we identified candidates for pS65Ub-dependent transcriptional upregulation by first identifying genes with reduced OA-dependent upregulation in PINK1 KO cells, then isolating those with promoter-dependent pS65Ub enrichment. 676 genes were upregulated on OA in WT relative to PINK1 KO cells of which 86 were pS65Ub-positive (**Extended Data Table 12**). RNA and ChIP Seq data for candidate pS65Ub-dependent genes with potential H2AK119Ub depletion are shown in **Figure 5F** and **Extended Data Figure 6F**. Of these candidate genes, BCL7A, which is implicated in neurodevelopment^91^, showed PINK1 activity dependent changes in protein levels (**Extended Data Figure 6F, G**).

These data suggest that pS65Ub is enriched at promoters of poorly expressed but dynamically regulated genes, poised for activation upon OA. More importantly, they are consistent with a novel axis of epigenetic regulation, in which mitochondrially-generated pS65Ub decorates chromatin and activates gene expression via depletion of H2AK119ub.

### Evidence for Functional pS65Ub-Dependent Polycomb Activation in Dopaminergic Neurons

Having identified pS65Ub as a potential activating epigenetic mark, we next investigated whether the decoration of chromatin with pS65Ub is sufficient to influence gene programs. We have mainly used mitochondrial toxins to model the chronic PINK1 activation associated with neurodegeneration. However, this approach makes it difficult to distinguish the effects of nuclear pS65Ub accumulation from those caused by mitochondrial damage or OMM-localized PINK1 activity. To address this, we developed a system that allows pS65Ub enrichment independently of mitochondrial injury or PINK1 recruitment. Specifically, we expressed a constitutively active *Tribolium castaneum* PINK1 variant for which the N-terminal mitochondrial targeting and transmembrane domains were replaced with three tandem NES tags, yielding a mitochondria- and nucleus-excluded cytoplasmic construct (iPink1). HeLa cells transiently expressing iPink1 phenocopied OA treatment by pS65Ub ChIP Seq analysis (**Extended Data Figure 7A, B**) without overt activity-dependent alterations in mitochondrial respiration when grown in basal media (**Extended Data Figure 7C**).

We used this system to determine whether pS65Ub enrichment drives transcriptional programs, focusing on polycomb targets. A temporal decrease in polycomb activity is causally linked with neuronal maturation, such that pharmacological inhibition of PRC2 activity accelerates neuronal maturation in vitro^37^. We therefore chose iPSC-derived dopaminergic neurogenesis as a model in which to study pS65Ub-dependent polycomb modulation. We transduced dopaminergic (DA) neuronal precursor cells (differentiation day 22) with lentiviruses encoding WT iPink1 and controls kinase dead (KD) iPink1 or BFP, each encoding EGFP connected by a T2A linker to track transduction rates. The cells were then differentiated to a mature state prior to analysis. Immunofluorescence analysis revealed that ∼30% of neurons were tyrosine hydroxylase-positive in optimization experiments, confirming the presence of DA neurons (**Extended Data Figure 7D**). Importantly, only WT iPink1 expression recapitulated OA-dependent nuclear pS65Ub enrichment and histone pS65Ubiquitination (**Figure 6A, Extended Data Figure 7E**). Due to the implication of pS65Ub in mitophagy and proteostasis^92^, we measured the relevant markers TIM23 and K48-linked ubiquitin, but found no obvious differences in expression, suggesting that any effects of WT iPink1 in the nucleus are specific (**Extended Data Figure 7E**).

**Figure 6.**
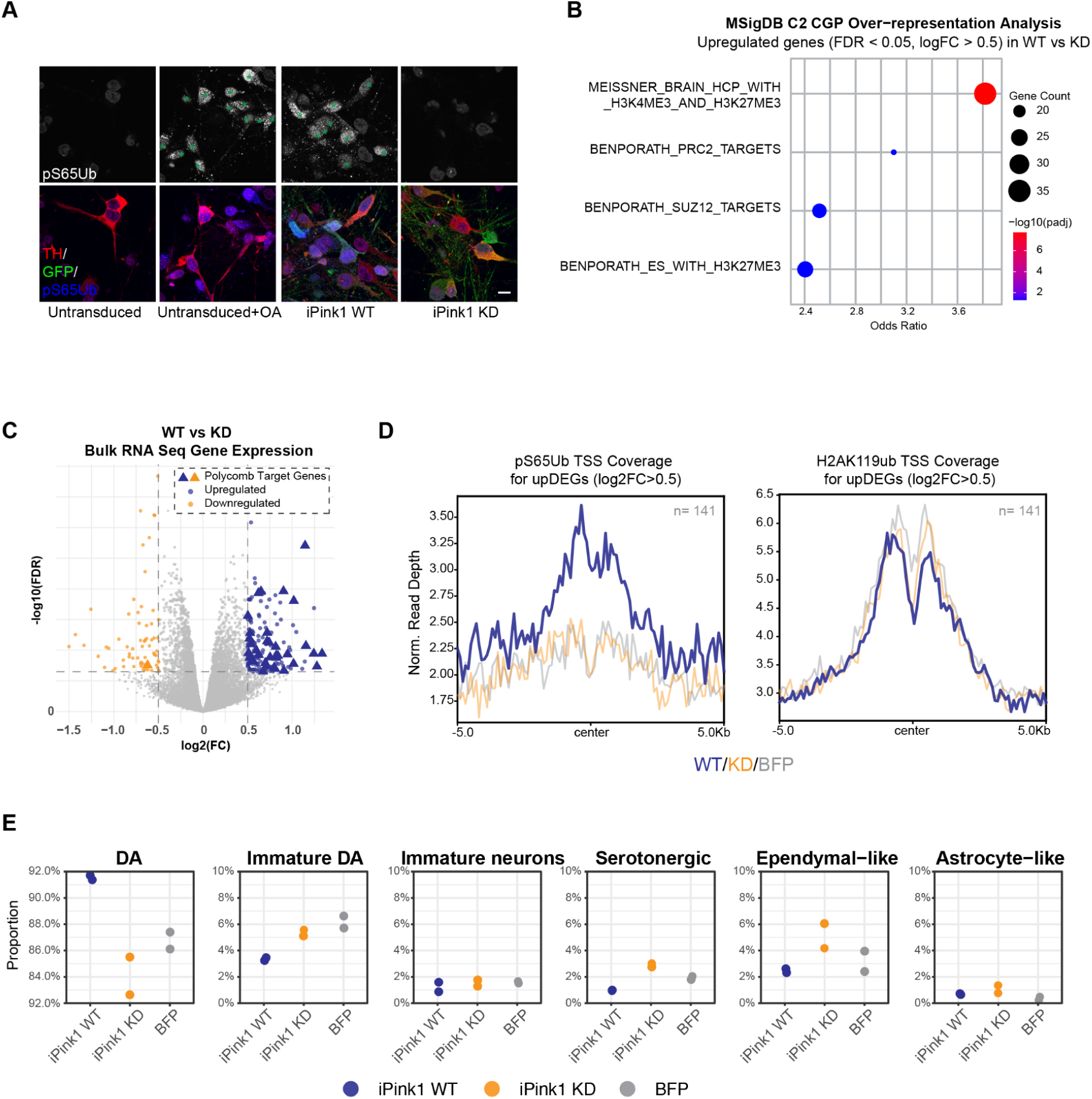
iPink1 overexpression promotes both polycomb gene expression and dopaminergic cell identity. **A** - iPSC-derived dopaminergic (DA) neuronal precursors were transduced with WT or kinase dead (KD) iPink1, or left untransduced. At maturity, untransduced cells were treated with OA (1µM, 24 hrs) or vehicle then the distributions of the DA marker tyrosine hydroxylase (TH, red), GFP (green) and pS65Ub (blue) were revealed by immunofluorescence, scale bar = 10µm. Cells nuclear pS65Ub enrichment are marked with green asterisks. **B** - DA neuronal precursors transduced with iPink1 WT or KD were analysed by bulk RNA Seq at maturity. An overrepresentation analysis was performed on genes significantly upregulated in WT vs KD transduced cells. **C** - Volcano plot showing significantly up and down-regulated genes (blue and orange respectively) in the WT vs KD comparison. Genes matching the top overrepresentation analysis term, ‘MEISSNER_BRAIN_HCP_WITH_H3K4ME3_AND_H3K27ME3’, were considered polycomb targets and are annotated as triangles. **D** - Metagene profiles showing averaged pS65Ub and H2AK119ub ChIP Seq signal at the transcription start site (TSS) of WT vs KD iPink1 upDEGs (Extended Data Table 13). **E** - Cell type proportions in iPink1 WT-, iPink1 KD- and BFP-transduced cell cultures defined by scRNA Seq gene expression signatures.

To capture pS65Ub-dependent phenotypic changes, we transduced cells as described above then, at maturity, performed parallel bulk RNA Seq and single cell RNA (scRNA) Seq analyses, as well as ChIP Seq with pS65Ub and H2AK119ub antibodies. Bulk RNA Seq analysis of all possible combinations (iPink1 WT vs KD, WT vs BFP, KD vs BFP, **Extended Data Table 13-15**), followed by overrepresentation analyses on the significantly up- and down-regulated genes, yielded significant terms in the WT vs KD upregulated gene set only. Strikingly, all four enriched terms were composed of polycomb target genes identified by ChIP Seq^93,94^ (**Figure 6B, C**), suggesting that WT iPink1 expression drives polycomb target gene upregulation.

To determine whether the expression changes were pS65Ub-dependent, we examined pS65Ub and H2AK119ub ChIP Seq signals at the promoters of iPink1 WT vs KD DEGs (**Extended Data Table 13**). WT-specific pS65Ub enrichment, along with evidence for H2AK119ub depletion, was detected in the upregulated gene set exclusively (**Figure 6D; Extended Data Figure 7F, G**), supporting the hypothesis that pS65Ub directly drives gene expression. Moreover, both bulk RNA Seq (overrepresentation analysis, **Figure 6B, C**) and ChIP Seq (H2AK119ub-enrichment, **Figure 6D, Extended Data Figure 7F, G**) implicate polycomb target genes as primary targets.

Finally, we performed scRNA Seq to resolve the cellular states at higher resolution and focus our analysis on DA neurons specifically. To isolate transduced cells for further analysis, we filtered for EGFP expression, revealing that in two independent replicates, only ∼14% of PINK1 WT or KD cells were exogene-positive (∼65% for BFP-transduced cells, **Extended Data Figure 7H**). While these proportions are likely underestimated due to a bias toward highly expressed genes, this heterogeneity likely underestimated the biologically relevant changes identified via bulk methods (RNA Seq and ChIP Seq).

Manual annotation of the main clusters using a combination of previously published and internal cell markers^95,96^ yielded six major cell types (**Extended Data Figure 7I, J**). Interestingly, WT iPink1 expression correlated with a proportional increase in mature DA neurons and a decrease in immature DA neurons (**Figure 6E**). To test whether polycomb gene expression might be implicated in the accelerated dopaminergic neuron maturation, we plotted the expression of brain-relevant polycomb target genes (i.e. the term most significantly overrepresented amongst genes upregulated on WT iPink1 expression (**Figure 6B**). Although not statistically significant, a modest WT iPink1-dependent increase in polycomb gene expression was observed exclusively in immature DA neurons (GSEA: normalized enriched score: 1.316; pval: 0.0008014 adjpval : 0.182505) and immature neuron subsets (**Extended Data Figure 7K**).

Collectively, these findings suggest that pS65Ub accelerates the maturation of DA neurons, potentially by promoting the expression of polycomb target genes.

### Nuclear pS65Ub Accumulates in the Diseased Brain

Our data show that nuclear pS65Ub accumulation correlates with PINK1 activity and reveal downstream effects of pS65Ub deposition on chromatin. We therefore asked whether nuclear pS65Ub enrichment is detected in more pathologically-relevant conditions, namely α-synuclein stress, a key pathological driver of PD implicated in PINK1 activation^97,98^, mitochondrial damage^25^, and epigenetic misregulation^27^. NGN2-cortical neurons were seeded with α-synuclein pre-formed fibrils, leading to the accumulation of phosphoserine 129 α-synuclein^99^, which correlates with pathological aggregation of endogenous α-synuclein (**Figure 7A; Extended Data Figure 8A**). Importantly, nuclear pS65Ub was elevated in cells undergoing α-synuclein stress (**Figure 7A**).

**Figure 7.**
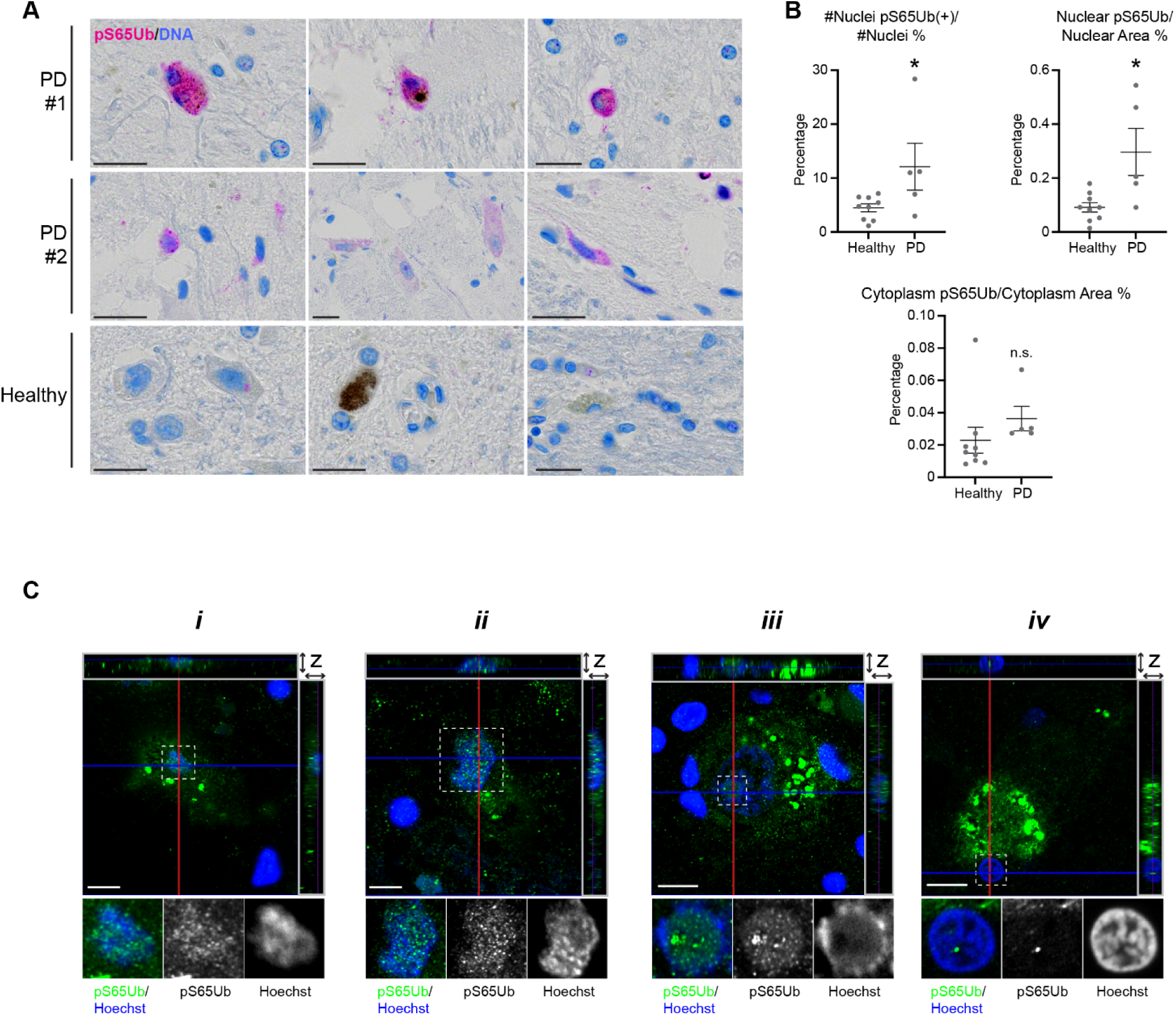
Disease-dependent accumulation of nuclear pS65Ub in the brain. **A** - SNpc section scans from five PD and nine healthy post-mortem brains were stained for pS65Ub (pink) and DNA (blue) before automated analysis of pS65Ub intensity. Representative images from individuals with and without PD are shown. **B** - The average percentages of nuclei containing pS65Ub signal and pS65Ub nuclear area vs total nuclear area (top), and pS65Ub cytoplasm area vs total cytoplasm area (bottom) were quantified in healthy and diseased tissues, 2-sample t-test. **C** - SNpc sections from clinical PD cases that are pathologically confirmed as Lewy body disease were stained with an independent pS65Ub antibody (Springer lab, in house), then Z-stack images were captured using super resolution microscopy. Multiple pS65Ub nuclear distributions were detected, with representative images shown. The largest images show representative images of single XY optical planes where pS65Ub is detected overlapping with the nucleus. Orthogonal view (‘z’, grey boxes) show Z-stack cross sections, revealing that the pS65Ub signal is nuclear. Nuclear pS65Ub displaying both diffuse (i, ii, iii) and punctate (ii, iii, iv) distributions were observed. Dashed boxes indicate areas magnified in inset, scale bar 10µm. *p<0.05, n.s. not significant.

We then examined pS65Ub distribution in diseased tissues. As PINK1 is mutated in only a minority of PD cases, we predicted that nuclear pS65Ub might accumulate in diseased tissues. To test this we assayed the distribution of pS65Ub in healthy and sporadic PD post-mortem brains by immunohistochemistry (IHC). To validate the IHC protocol, we confirmed that phosphatase-sensitive pS65Ub signal was detected in HeLa cell and in PD substantia nigra pars compacta (SNpc) samples (**Extended Data Figure 8B, C**). In sections containing the SNpc, pS65Ub was detected in both neurons and glia with cytoplasmic distributions similar to previous reports^57,98^ (**Extended Data Figure 8D**), however we also observed signal that overlapped with nuclei (**Figure 7A**). Sections stained with an independent pS65Ub antibody yielded similar pS65Ub distributions further supporting the specificity of the signal (**Extended Data Figure 8E**).

Next, we stained a range of age-matched PD and healthy SNpc samples using the pS65Ub IHC protocol to assess nuclear pS65Ub prevalence in human disease (**Figure 7A**; **Extended Data Figure 8F**). Unbiased image analysis revealed that the fraction of nuclei positive for pS65Ub, and both nuclear and cytoplasmic pS65Ub signal was increased in a subset of PD subjects compared to healthy controls (**Figure 7B**). Finally, we turned to super resolution microscopy to more confidently localize pS65Ub within nuclei in tissues. We stained SNpc sections from post-mortem brains of clinical PD cases (pathologically confirmed as Lewy body disease) using an independent pS65Ub antibody which has been widely used for IHC in tissues^57,98^. We then acquired super resolution Z-stack images, identifying intranuclear pS65Ub in both punctate and diffuse distributions (**Figure 7C**), corroborating the chromogenic data in **Figure 7A**. These data suggest that accumulation of nuclear pS65Ub is associated with Parkinson’s disease.

## Discussion

The coordination of mitochondrial health and nuclear gene expression is increasingly recognized as a relevant factor in neurodegenerative disorders^31,33,100–102^. In this study, we show that the primary product of PINK1, pS65Ub, accumulates in the nucleus in response to mitochondrial damage where it functionally alters gene expression, implicating pS65Ub in mitochondria to nucleus signaling (**Extended Data Figure 9**). Importantly, we observed nuclear pS65Ub accumulation in PD brain samples, suggesting that nuclear pS65Ub signaling may occur *in vivo*.

The accumulation of pS65Ub in the nucleus and on chromatin adds a new axis of communication between mitochondria and nuclei, one with unique and intriguing features. Firstly, our data support a secondary messenger like system, whereby pS65Ub is released from damaged mitochondria to affect changes in the nucleus. Nuclear pS65 ubiquitination corresponds with PINK1 activation at mitochondria, being detected in minutes then peaking at 4-8 hrs following CCCP-dependent mitochondrial membrane potential collapse, in keeping with previously reported dynamics for CCCP-dependent PINK1 stabilization^103^. Our data suggest that Parkin confines pS65Ub at mitochondria, likely via enhanced ubiquitin polymerisation at the OMM or via autophagic capture of both pS65Ub and PINK1 in mitolysosomes. This adds to a stream of Parkin-independent PINK1 phenotypes^104^, including modulating of oxidative phosphorylation efficiency^105^, promoting the cytoprotective functions of TRAP1^106^, providing a ‘molecular memory’ to prime response to sequential mitochondrial insults^77^, and pertinently, impairing proteasomal activity via excess pS65Ub generation in the context of neurodegeneration^107^. The relevance of these mechanisms in disease is an important avenue for further study.

Secondly, we provide evidence for a new mode of pS65 ubiquitination. In the canonical pS65 ubiquitination pathway, pS65Ub activates Parkin at the OMM, which then synthesizes polyubiquitin chains without incorporating pS65Ub directly^20,81–83^. This generates more substrates for PINK1 in a feed-forward loop, where pS65Ub-dependent inhibition of the mitochondrial DUB USP30 may enhance signal further^15,19–21^. With noncanonical pS65 ubiquitination however, pS65Ub is directly ligated by the E3 ligases. Furthermore, for H2AK119, our data show that pS65Ub reduces E3 activity as a suboptimal substrate, while potentiating deubiquitinases, ultimately counteracting substrate ubiquitination.

Thirdly, we show that pS65Ub itself is attached to chromatin as a novel epigenetic mark. Multiple strands of evidence point to H2AK119 being a major pS65Ub acceptor on chromatin. While our data suggest that a small fraction of the total histone pool is pS65 ubiquitinated, simultaneous E3 ligase inhibition and DUB potentiation may enhance pS65Ub’s ability to drive H2AK119 deubiquitination (particularly at USP16 and USP21 target genes). It is possible that pS65Ub-dependent gene activation is most apparent at genes that are otherwise poised for activation, i.e. needing only a small ‘push’. Supporting this notion, analysis of the pS65Ub-dependent genes upregulated in dopaminergic neurons revealed that the most enriched term corresponded to ‘bivalent genes’ (genes that are repressed by polycomb marks at their promoters while remaining poised for expression due to the presence of activating marks such as H3K4me3) identified in whole-brain lysates^94^. Of potential relevance, recent work shows that loss of H2AK119ub at a subcluster of bivalent promoters precedes and likely facilitates increased gene expression in Huntington’s disease mouse models^108^.

Lastly, the absence of nuclear-encoded mitochondrial genes from the pS65Ub gene set, together with a subtle reduction in mitochondrial respiration on iPink1 expression, suggests that extra-mitochondrial pS65Ub might not promote mitochondrial homeostasis, although this warrants further study. Instead, our data support a model in which pS65Ub drives polycomb target gene de-repression.

Whether pS65Ub-dependent polycomb de-repression is protective or deleterious may depend on the magnitude and duration of stress. It could support neuronal identity or differentiation, consistent with links between *PINK1* and other PD risk genes with defects in dopaminergic neuron differentiation^109–111^. Alternatively, chronic pS65Ub-dependent H2AK119 deubiquitination may destabilize Polycomb domains^39,45–47^, inadvertently relieving gene repression as seen in mouse brains^55,112^, where genetic ablation of PRC2 activity is causally linked to neurodegeneration^55^. Notably, age-dependent de-repression of polycomb target genes is observed in the human brain^112^.

We consider this latter ‘toxic gain of function’ model to be particularly compelling, as a gradual loss of proper epigenetic information (known as ‘epigenomic erosion’)^108,113–116^ is emerging as a common factor in neurodegenerative diseases. Even more pertinently, recent work identifies PRC1- and H2AK119ub-dependent de-repression of developmental genes as a driver of epigenomic erosion in Huntington’s disease^108^. Together with our findings, this raises the possibility that chronic pS65Ub exposure in adult neurons promotes epigenomic erosion through Polycomb domain destabilization, an important avenue for future investigation.

The prominence of the pS65Ub-histone band on Western blots together with proteomic and genomic evidence pointing towards H2AK119ub encouraged us to more narrowly focus on this as the major mono pS65Ub acceptor. Our analytical strategy was therefore constrained by technical challenges in isolating the H2AK119-pS65Ub pool, including the lack of a specific antibody and the pleiotropic effects of genetic perturbations. For example, RING1A/B double knockout abolishes H2AK119 pS65 ubiquitination, but also eliminates global H2A ubiquitination and disrupts the non-catalytic functions of the PRC1 complex^38^, while redistributing pS65Ub to alternative acceptors. Similar limitations applied to assessing nuclear pS65Ub in mitophagy, as available tools perturb both pathways. Developing approaches to selectively isolate discrete pS65Ub pools therefore remains an important goal for future studies.

Taken together, we propose a novel secondary messenger-like mechanism whereby mitochondrially-generated pS65Ub dynamically regulates epigenetic signaling in the nucleus. After generation at the surface of damaged mitochondria, pS65Ub likely diffuses into the nucleus, where it is ligated to substrates by at least one resident E3 ligase, a process termed noncanonical pS65 ubiquitination. In the nucleus, it becomes enriched at promoters of developmental genes as a novel epigenetic mark, H2AK119pS65Ub. pS65Ub reciprocally regulates E3 ligases and deubiquitinases, biasing H2A towards its K119 unmodified state and driving functional gene programs via polycomb target de-repression (**Extended Data Figure 9**). The enrichment of nuclear pS65Ub in diseased brains indicates that the mechanisms deciphered here may also be active in PD. We speculate that excess nuclear pS65Ub could rewire polycomb signaling and contribute to epigenomic erosion during neurodegeneration.

## Materials and Methods

### Cell Culture

HeLa and HEK293T cells were cultured in high glucose DMEM, and NCI-H226, murine melanoma and murine PDAC cells were cultured with RPMI, both supplemented with 10% FBS, 1X GlutaMAX Supplement and 1X Pen Strep. Cells were grown in a humidified incubator maintained at a 37°C with 5% CO2. Cells were transfected when 60-80% confluent and analysed when confluence had reached ∼90% . Lipofectamine 2000 was used for all transfections, used as per the manufacturer’s instructions. Plasmids were transfected the day after seeding at 0.4-1.5µg/ml media then analyzed 1 day later. For knock down experiments siRNA was reverse transfected on day 1 and again on day 2, both at 50 nM final concentration, before analysis on day 4.

The human induced pluripotent stem cell (hiPSC) line, iP11N, stably expressing doxycycline- inducible Neurogenin-2 (NGN2) cassette, was used in Figures 1 and 2. The NGN2 iNeuron differentiation and maturation was performed following the protocol previously published with some modifications^117^. In brief, hiPSC were maintained in mTeSR™ Plus media using the substrate iMatrix-511. When iPSC cultures reached ∼75% confluency, they were dissociated using ReLeSR™. On day 0, ∼75% confluent iPSC cultures were dissociated into single cells using Accutase and seeded at a density of 40,000 cells / cm 2 on iMatrix-511 substrate in induction media consisting of DMEM/F-12, B27, N-2 Max, GlutaMax, NEAA, 10 µM SB431542, 2 µM XAV939, 10 µM DAPT, 100 ng/ml Noggin and 1 day use of Y27632. Neural induction was achieved by supplementing induction media with 3 µg/ml doxycycline and performing daily media changes for 5 consecutive days. On day 5, the media was changed to maturation media consisting of Neurobasal medium, B27, GlutaMax, 20 ng/ml BDNF, 20 ng/ml GDNF, 1 ng/ml TGFβ3, 500 µM cAMP, 10 µM DAPT) with doxycycline concentrations reduced to 2 µg/ml. Maturation media was also supplemented with Cytosine β-D-arabinofuranoside hydrochloride at 5 µM (to eliminate any remaining dividing cells) for 2 days. On day 7, cultures were treated with Accutase for 20 minutes and then quenched with maturation media containing Deoxyribonuclease I. Cultures were gently triturated using a serological pipette, passed through a 40 µm cell strainer, and centrifuged (300g for 5 minutes). Pellets were resuspended in CryoStor cryopreservation media, and banks of day 7 induced NGN2 (iNGN2) neurons were created. Day 7 iNGN2 neurons were then thawed and seeded in maturation media supplemented with 2 µg/ml doxycycline on Poly-D-lysine and iMatrix-511-coated tissue culture plates. Doxycycline treatment was continued for the next 7 days with ½ media exchanges every other day to day 7. Following the 7-day doxycycline treatment, ⅓ media exchanges, using maturation media, were performed 3 times per week for the duration of the experiment.

Dopaminergic neurons were differentiated based on previous methods^118,119^. In brief, hiPSC were maintained in E8 using the substrate Geltrex. When iPSC cultures reached ∼75% confluency, they were dissociated using ReLeSR™. On day 0, ∼75% confluent iPSC cultures were dissociated into single cells using Accutase and seeded at a density of 400,000 cells / cm2 on iMatrix-511 substrate in Neurobasal medium supplemented with B-27, N-2 Supplement B, GlutaMax, 10 uM SB431542, 250 nM LDN, 500 ng / mL SHH, 100 ng / mL Noggin, 0.7 uM CHIR99021 and Y27632. Complete media changes were performed daily (without Y2763) for 3 days following. From days 4-6, CHIR99021 concentration was increased to 7.5 uM, again with complete media changes daily. From days 7-9, SB431542, LDN, SHH and Noggin were removed and 7.5 uM CHIR99021 was maintained for the daily media changes. On day 10 media was fully changed to a Neurobasal base media (NB) consisting of Neurobasal medium supplemented with B-27, GlutaMax, 200 uM Ascorbic Acid, 0.5 mM Dibutyryl cyclic-AMP sodium salt, 20 ng / mL hBDNF, 20 ng / mL hGDNF, and 1 ng / mL hTGFβ3. On day 11, cultures were dissociated into single cells using Accutase and replated into iMatrix-511 coated vessels at a seeding density of 700,000 cells / cm2 using NB media containing Y2763. Daily full media changes were performed up to day 15 using NB (without Y2763). The cultures were again replated on day 16 by dissociating with Accutase and seeding into iMatrix-511 coated vessels, using NB media containing 10 uM DAPT and Y2763 at 700,000 cells / cm2. From days 17-20, full media changes were performed daily using NB media containing DAPT. On day 21 cultures were again dissociated into single cells, this time using Papaine and replated into Poly-D-Lysine and iMatrix-511 coated vessels at a seeding density of 500,000 cells / cm2 using NB media containing DAPT and Y2763.

Control (#SFC-840-03, #SFC-841-03, #SFC-856-03), PINK1 (Q456X/Q456X, #SFC-824; Q456X/Q456X, #SFC-825; Q456X/Q456X, #SFC-826)^120^ and PRKN (c1072tdel/Ex7del, #SFC-817); R275W/Ex4 del, #SFC-818; R275T/C352R, #SFC-821)^121^ PD patient-derived iPSCs^120^ were differentiated to dopaminergic or NGN2-cortical neurons using these methods with minor modifications^120,122^, and seeded with α-synuclein pre-formed fibrils as previously described^99^. Participants gave signed informed consent to mutation screening and derivation of iPSC lines from skin biopsies (Ethics committee: National Health Service, Health Research Authority, NRES Committee South Central, Berkshire, UK, REC 10/H0505/71).

Control primary fibroblasts (#106-05A) were from Cell Applications, Inc., PINK1 (Q456X/Q456X, #sc1028) and Parkin (Ex4-7 del / c.203_204 del AG, #sc1064) mutant fibroblasts are available from the NINDS stem cell repository. Fibroblasts were cultured in DMEM containing 10% FBS. Fibroblast medium was supplemented with 1% non-essential amino acids and 1% Penicillin/Streptomycin.

Post transfection, cells were incubated with doxycycline (1µg/ml) for 20 hrs to induce USP16 or USP21 expression. CCCP, OA, Valinomycin, Rotenone, Paraquat, PRT062607, MB-10, MG132 and Rapalog treatments were performed as described in the figure legend. Where appropriate, DMSO was used as a vehicle control.

### Cell Line Generation

To generate RING1A/B knockout lines, cells were co-transfected with a plasmid expressing Cas9 + blasticidin resistance and a plasmid expressing puromycin resistance and either a *RING1B* or *luciferase* gRNA. Double transfected cells were selected using blasticidin and puromycin for 4 days before expansion of surviving cells. After validation of RING1B knockout status, these cells then underwent another round of CRISPR KO with gRNAs targeting RING1A. The guide RNAs used were: *Luciferase*: 5’CCG GCG CCA TTC TAT CCG C. For human cell lines: *RING1A*: 5’AAG ATC TAT CCT AGC CGG G; *RING1B*: 5’GAG TTA CAA CGA ACA CCT C. For mouse cell lines: *Ring1a*: 5’ATC TCT GTA CCA TCC ATG A; *Ring1b*: 5’GTG CAT CAA AGT TCG GGT C.

To generate iPink1 overexpressing HeLa cells, we engineered *Tribolium castaneum* Pink1 with the N-terminal 127 amino acids (encoding the mitochondrial target sequence and transmembrane domain) replaced with three tandem HIV Rev protein nuclear export signal sequences and encoding a C-terminal V5 tag, then inserted wild type or kinase dead (encoding K196A, D337A, D359A mutations) NES-iPink1-V5 into PiggyBac transposon vectors. The PiggyBac vectors were transfected into HeLa cells with a mixture of transposon:transposase plasmids at 2:1 as described above. After 3 days, cells were selected with blasticidin, then doxycycline-induced iPink1 expression was confirmed by immunofluorescence analysis.

### Formalin-Fixed Paraffin-Embedded (FFPE) Samples

For control samples, HeLa cell lines were treated with CCCP for 1 hr or either CCCP or DMSO for 4 hrs. Treated cell lines were pelleted then made into FFPE cell line microarray blocks. For experimental samples, archival FFPE tissue blocks of the SNpc of five idiopathic PD and nine control cases, male and female ages 64-85 were acquired. All samples were cut into four micron thick sections then mounted on charged microscope slides.

### Chromogenic Immunohistochemistry

Sections were baked at 70°C for 30 mins, then deparaffinized in xylene and rehydrated through graded alcohols to distilled water, then pretreated with 1x Target Retrieval solution (Citrate pH6.1) for 20 mins at 99°C, and rinsed in dH2O. Single, chromogenic immunohistochemistry staining was performed on the Ventana Discovery Ultra autostainer (Roche Tissue Diagnostics, Tucson, AZ, USA).

Anti-pS65Ub antibody (Cell Signaling Technology, #62802) was diluted in 3% bovine serum albumin in PBS to 0.78ug/mL. Sections were incubated for 60 mins at room temperature, followed by Ventana Discovery Anti-Rabbit HQ for 16 mins at 37°C, then Ventana Discovery Anti-HQ HRP for 16 mins was used as the detection system. Ventana Discovery Purple Kit was incubated for 12 mins for antibody visualization. Ventana Hematoxylin II was used to counterstain. For the additional pS65Ub antibody clone tested, anti-pS65Ub (Cell Signaling Technology, #70973) was used at 1ug/ml with the same IF protocol as clone #62802, at 0.5-2.5ug/ml with Omnimap anti-Rabbit HRP detection. To determine the specificity of the chromogenic immunohistochemistry assays, we confirmed that pS65Ub signal was reduced by lambda phosphatase treatment, and stained parallel samples with Rabbit mAb IgG Isotype Control. For lambda phosphatase assays, samples were deparaffinized, underwent Target Retrieval, then treated with lambda phosphatase at 8,000 Units/mL for 60 mins at 37°C, followed by IHC staining for pS65Ub

### Immunohistochemistry Image Analysis

Immunohistochemically stained slides were imaged in brightfield mode on the Olympus Slideview VS200 scanner (Olympus Scientific, Waltham, MA). Images were acquired using the 40x silicone oil immersion objective (NA 1.4) at a resolution of 0.137 microns per pixel. Image processing was performed using morphological operations and color thresholding in MATLAB (MathWorks, Natick, MA). The ’2D_versatile_he’ pre-trained model in StarDist^123^, a deep learning based objection detection python library, was used for nuclei segmentation. pS65-Ub positive staining was detected using the machine learning pixel classification module of ilastik^124^. The random forest based pixel classifier was trained using manual labels in interactive mode. The nuclei and pS65-Ub masks were dilated by 5 microns to account for the cytoplasmic space. The number of nuclei that contained positive pS65-Ub staining was normalized to the total nuclei in the tissue sample. Nuclear and cytoplasmic pS65-Ub positive area was normalized to the nuclear and cytoplasmic area, respectively. All image analysis was performed in a blinded manner and results were reviewed by a neuropathologist.

### Microscopy

Cells were seeded on 96 well imaging plates (Revvity, #6005225) or on Poly-D-lysine-coated coverslips before fixation in PBS 4% paraformaldehyde (PFA, 20 mins, room temperature/RT). Cells were permeabilized (PBS + 0.2% Triton X-100, 5 mins RT) then blocked (PBS + 5% BSA, 30 mins RT), then incubated with primary then secondary antibodies diluted appropriately in PBS + 5% BSA. Where indicated, cells were incubated with Hoechst stain (NucBlue, Thermo) alongside secondary antibodies to stain nuclei. Cells grown on coverslips were mounted on slides before imaging. Imaging was performed on a Leica SP8 inverted confocal microscope with a 63X objective lens, with data processed using Adobe Photoshop or FIJI.

For experiments using iPSC-derived NGN2-cortical and dopaminergic neurons (excluding α-synuclein seeding and PINK1/Parkin mutant experiments, described below), cells were grown on 96-well Phenoplates (Revvity) were treated with either OA or vehicle control for 6 or 24 hrs at 1µM. Following treatment, neurons were fixed with PBS 4% PFA for 15 minutes at 37°C. Fixed neurons were then washed three times with PBS and permeabilized with 0.2% Triton X-100 in PBS for 10 minutes, followed by blocking in 5% normal donkey serum (Abcam Inc., #ab7475) and 1% BSA (Millipore, #A7979) in PBS for 1 hr at room temperature. Primary antibodies against tyrosine hydroxylase, pS65Ub and HSP60 were applied overnight at 4°C in blocking solution. After three washes with PBS, secondary antibodies, Alexa Fluor-conjugated secondary antibodies diluted in blocking buffer were applied for 1 hr at room temperature. Nuclei were counterstained with Hoechst (NucBlue, Thermo) during the secondary antibody incubation. Cells were kept in the final PBS wash prior to imaging. Images were acquired using a Zeiss LSM980 Confocal microscope equipped with 639 nm, 561 nm, and 405 nm lasers and a Plan-Apochromat 20x/0.8 M27 objective. Z-stack images were captured with a step size of 0.4275 μm, totaling 20 slices and a total thickness of 8.55 μm, with an 8-bit depth. Maximum intensity projections of the z-stacks were generated using FIJI software.

For analysis of PINK1/Parkin mutant dopaminergic and α-synuclein-seeded iPSC-derived NGN2-cortical neuron cultures, cells were imaged in 96-well format PhenoVue plates, as previously described^122^. Briefly, Cells were washed, fixed with 4% paraformaldehyde solution for 20 min and then washed twice. Fixed cells were incubated with permeabilisation buffer (10% v/v FBS, in PBS) for 15 min, and blocked for 1 h at room temperature in 0.1% v/v Triton-X,10% v/v normal goat serum, in PBS. Cells were incubated with primary antibodies (monoclonal anti-pS129 alpha-synuclein; polyclonal anti-pS65Ub (Millipore, ABS1513-I-AF488, for NGN2 neurons); monoclonal anti-pS65Ub (Cell Signalling, 62802, for dopaminergic neurons); polyclonal anti-MAP2 (Merck, ab5543) diluted in blocking solution at 4°C overnight. Cells were washed and incubated with secondary antibodies and NucBlue (Thermo), and were diluted in blocking solution for 1 h. Cells were then washed and imaged using an Opera Phenix High-Content Imaging system (Revvity). Some images used in the analysis presented in Figure 1B have been previously analysed for a-synuclein marker pS129 in the following manuscript^99^.

For live cell imaging experiments, HeLa cells were seeded on PhenoPlate 96-well plates (Revvity, #6055302) then allowed to adhere overnight. The next day cells were treated with CCCP in basal growth medium (DMEM) at defined concentrations for 3 hours. At this point, cells were incubated in DMEM containing the specific concentration of CCCP along with MitoTracker™ Green FM (100nM) and MitoProbe™ TMRM at 20nM and NucBlue reagent at 1:100 v/v. After 30 mins incubation, the dye-containing medium was removed and treatment medium (DMEM + CCCP), and cells were taken for live cell imaging, which was completed ∼ 4 hrs after the initial treatment began. Images were acquired using a Leica Thunder widefield microscope with DAPI, FITC and Texas Red filter sets at 20x magnification operated with Leica LAS X software. Ten regions were imaged per condition. For analysis, the machine learning program Ilastik was used to identify and segment Mitotracker Green and MitoTMRM areas^124^. Ilastik segmentations were imported into Fiji and the area of identified segmentations was measured using “Analyze particles”^125^. MitoTMRM area/Mitotracker GREEN area was calculated across all conditions, and the ratio of each region imaged was normalized to the average of the 0 µM condition and plotted in GraphPad PRISM.

For high content imaging of fibroblasts, 100 µl of a 50K/ml cell suspension was plated into 96-well plates (Greiner, #655090) and cells allowed to attach overnight. Fibroblasts were treated with 1µM valinomycin, fixed with 4%PFA, permeabilized with 1% Triton for 15 min and blocked with 10% normal goat serum. Cells were stained with antibodies against pS65Ub (in-house^57^; Cell Signaling Technology, #62802; Cell Signaling Technology, #37642; MilliporeSigma, ABS1513, Signaling Technology, #70973) and TOM20, followed by incubation with appropriate secondary antibodies. Nuclei were stained with Hoechst 33342. Cells were imaged on a BD pathway 855 (BD Biosciences) using a 20x objective and a 2x2 montage (no gaps) with laser autofocus. Quantification of pS65Ub signal in the cytoplasm and nucleus were performed using regions of interest that were defined based on the Hoechst staining.

Immunofluorescence staining of post-mortem human FFPE brain tissues was performed as previously described^98^ . Sections were cut at a thickness of 5µm and mounted onto positively charged slides to dry overnight at 60°C. After sections were deparaffinized and rehydrated, antigen retrieval was performed by 30-minute steaming in deionized water. Following 1 hr blocking with the serum-free Protein Block, sections were incubated overnight at 4°C with primary antibody against pS65-Ub (in-house^57^) diluted in Dako Antibody Diluent. The next day sections were incubated with secondary antibody and DAPI at room temperature for 1.5 hrs, followed by 3 mins incubation in Sudan Black and then coverslipped in fluorescence mounting medium (Dako, S3023). After immunofluorescence staining, super-resolution confocal (Airyscan) images were taken with a LSM 880 microscope (Zeiss, Oberkochen, Germany) with Z-stacks.

### Western Blotting

Cells were lysed on ice in HNTE (20mM HEPES pH7.5, 150mM NaCl, 5mM EDTA, 1% w/v Triton X-100, 1X HALT Protease and Phosphatase Inhibitor Cocktail), then transferred to 1.5ml Eppendorf tubes before water bath sonication to solubilize chromosomal DNA. Whole-cell lysates were mixed with LDS sample buffer and reducing agent to 1X then incubated at 95°C for 5 mins before protein separation using NuPAGE Bis-Tris Mini Protein Gels, 4–12% (Thermo). Proteins were transferred onto nitrocellulose using the iBlot 2 Dry Blotting System (Thermo) and blot sections incubated in either PBST or TBST with 5% skim milk powder (Neogen) or 3% fraction V BSA (Rockland) respectively for 1 hr. Blots were incubated with appropriate primary and secondary antibodies before HRP activation using ECL Select buffers (Cytiva) and digital imaging on iBright 1500 (Thermo). Image J (NIH) was used for quantification.

### Subcellular Fractionation

Crude subcellular fractionation was performed with the Cell Fractionation Kit (Cell Signaling Technology) using CIB, MIB and CyNIB buffers as per manufacturers instructions. To generate the Tx100 washed nuclear fraction in Figure 3C, samples were split in half after incubation with MIB buffer. One half was processed as described in the datasheet to generate membrane-bound organelle and nuclear pellets. The second half was diluted 15 fold in HNTE buffer, vortexed for 15 seconds, incubated on ice for 5 mins then centrifuged at 21,130xG for 5 mins, 4°C. Crude and washed nuclear pellets were water bath sonicated to solubilise genomic DNA then prepared for Western blot analysis as described above.

To generate nuclear lysates for anti-pS65Ub affinity purification mass spectrometry (pS65Ub APMS) experiments, PINK1 WT and KO HEK293T cells were treated with either DMSO or OA for 18 hrs then crude nuclear fractions were generated using the Cell Fractionation Kit (Cell Signaling), with all fractionation buffers supplemented with DUB inhibitor PR-619 (100µM). Nuclear pellets were mixed with Urea Lysis Buffer (8M Urea, 20mM HEPES pH7.5, 135mM NaCl, 1% Triton X-100, 1.5mM MgCl2, 5mM EDTA, 1X HALT, 10% glycerol and 100 μM PR-619) then water bath sonicated. Lysates were then diluted with lysis buffer constituents without urea to bring urea concentration down to 4M for immunoprecipitation.

To generate nuclear lysates for nucleosome affinity purification mass spectrometry (nucleosome APMS) experiments, we modified a previously published protocol^126^. Four 15cm dishes were seeded with HeLa cells and treated with OA for 18 hrs when fully confluent. Cells were detached using trypsin and washed with full media followed by PBS, then cell pellets were resuspended in 4ml nuc isolation buffer (10 mM HEPES–KOH pH 7.9, 5 mM MgCl2, 0.25M Sucrose, 0.1% NP40 to, 1mM DTT 1X HALT and 100µM PR-619), vortexed for 15 seconds then incubated on ice for 10 mins. Nuclei were pelleted at 9800xG for 10 mins, 4°C. To extract nuclear proteins, nuclei were resuspended by gentle pipetting in extraction buffer (20 mM HEPES–KOH pH 7.9, 420mM NaCl, 25% glycerol, 1.5 mM MgCl2, 0.2 mM EDTA, 0.1% NP-40, 1 mM DTT, 1X HALT and 100µM PR-619), then incubated rotating at 4°C for 30 mins. Nuclei were spun at 16,000g for 10 mins in a 4°C centrifuge and the supernatant was transferred to a fresh 1.5ml Eppendorf tube before briefly sonicating as a precaution to clear contaminating DNA. The nuclear lysate was diluted to 100mM NaCl in nuc isolation buffer containing 0mM NaCl and aliquots were flash frozen and stored prior to use at -80°C.

### Immunoprecipitation

For pS65Ub APMS experiments, nuclear lysates were prepared as described above. The protocol followed was previously reported^58^, with the following modifications: 12.5 μg anti-pS65Ub antibody was bound to 50 μL Protein G Dynabeads, and immunoprecipitations performed with ∼800μg pre-cleared lysate. Beads were washed 3 times with 4M Urea Lysis Buffer, then eluted in by 5 mins shaking at RT in 2x50µl 0.15% TFA. Eluate pH was altered to 8.5 using HEPES buffer, samples were flash frozen and kept at -80°C before analysis by mass spectrometry at a later date.

For nucleosome APMS experiments, unmodified, monoubiquitinated and pS65 ubiquitinated biotinylated mononucleosomes (Epicypher), or reaction mixtures without nucleosomes, were generated as bait samples (see ‘*In Vitro* Ubiquitination and Deubiquitination Assays’ below). To immobilize nucleosomes, 200µl reaction mixtures were diluted 1:4 with 0mM NaCl nuc isolation buffer, before rotation for 1.5 hrs at 4°C with 50µg washed streptavidin magnetic beads per 1µg biotinylated nucleosomes. The bead-bound nucleosomes were washed once with 1ml 50mM NaCl nuc isolation buffer and once with 1ml 100mM NaCl nuc isolation buffer. For prey samples, frozen nuclear lysates aliquots from 7 independent experiments (described above) were thawed, pooled then precleared by rotating at 4°C for 90 mins with 5µg streptavidin beads per 1µg lysate. The precleared lysate was then split into 4x∼275µg aliquots then used to resuspend the bead-bound nucleosomes, which were rotated at 4 hrs at 4°C. The pulldown samples were then gently washed with 3x1ml 100mM NaCl nuc isolation buffer before elution in 2x75µl 0.15% before neutralisation with NaOH and flash freezing. Samples were stored at -80°C before mass spectrometry analysis.

For *in vitro* deubiquitination assays, HEK293T in 10cm dish format were transfected with 3xHA-USP16, 3xHA-USP21 or empty vector. Post-transfection, cells were refreshed with culture media containing doxycycline to induce protein expression (see ‘Cell Culture’ section). Cells were lysed in 750µl HNT buffer (20 mM HEPES pH7.5, 150 mM NaCl, 1% w/v Triton X-100, 1X HALT Protease and Phosphatase Inhibitor Cocktail). The lysates were centrifuged at 21,130xG (5 mins, 4°C) and the supernatant transferred to a fresh 1.5ml Eppendorf. Post-nuclear supernatants were rotated with 62.5µg anti-HA magnetic beads at 4°C for 4 hrs, then washed with 3x1ml 150mM NaCl HNT and 1x1ml 50mM NaCl HNT.

### *In Vitro* Ubiquitination and Deubiquitination Assays

RING1B-PCGF1-5 proteins were kindly provided by Andrea Cochran and were previously reported^127^. When comparing the relative activities of RING1B-PCGF1-5, *in vitro* ubiquitination assays were performed as described previously^127^ with the following modifications: Reaction mixtures (50nM E1 (UBE1), 50nM E2 (UBE2D3), 300 nM RING1B–PCGF, 20 μM Ub (either WT or pS65Ub) and 150 nM mononucleosomes diluted in 50 mM HEPES pH 7.2, 0.2 mM DTT, 5 mM MgCl2, 2.5 mM ATP and 2 mg/ml BSA) were incubated at room temperature for 1 hr. Enzymatic reactions were stopped by addition of LDS and reducing agent to 1X dilution.

For nucleosome APMS, *in vitro* ubiquitination assays were performed as above, however only one E3 was used (RING1B-PCGF1). Additionally, to balance modification rates between Ub types while minimizing H2AK119 multimonoubiquitination, differing E3 concentrations were used in WT (25.5nM) and pS65Ub (72nM) mixtures. Reactions were stopped by washing as described above.

For *in vitro* deubiquitination assays we followed a modified version of Sahtoe et al^128^. We used the same E3 ratios as for nucleosome APMS experiments, but performed reaction in Modified E3 Buffer (50 mM HEPES pH 7.2, 0.2 mM DTT, 5 mM MgCl2, 2.5 mM ATP, 2 mg/ml BSA, 2µM ZnCl2 and 50mM NaCl). After incubating 1 hr at room temperature, reactions were stopped by addition of apyrase. We then washed DUB-bound beads (see ‘Immunoprecipitation’ section above) in Modified E3 Buffer, and split each sample into two equal aliquots, to which an equal amount of either the WT or pS65 ubiquitinated nucleosomes were added. Samples were incubated at 30°C for 0-60 mins with gentle resuspension every 10 mins. Aliquots were removed and mixed with LDS and reducing agent to stop the reaction at indicated times.

### Histone Purification

Histones were purified from fully confluent 10cm dishes using the Histone Purification Mini Kit (Active Motif, #40026) according to manufacturers instructions.

### Electroporation

The introduction of pS65Ub or HA-ubiquitin into PINK1 knockout HeLa cells via electroporation was performed using the Neon Transfection System 10 μL Kit (Thermo, #MPK1025) as per the manufacturers instructions. Specific details follow: The electroporation settings were Pulse Voltage: 1005v; Pulse Width: 35; Pulse Number: 2. 2 aliquots of washed resuspended PINK1 KO HeLa cells (2x 1.25x10^6^) were pelletted then resuspended in either 100µl Buffer R + 25µl 250µM pS65Ub or 100µl Buffer R + 25µl 250µM HA-Ub, 10x10µl from each mixture was electroporated, then all cells transfected with same protein to combined in a 1.5ml Eppendorf tubes containing 500µl DMEM culture media without Pen Strep. The combined mixtures were then split in 3 and diluted into further 500µl DMEM aliquots which were incubated at 37°C. After 15, 60 and 120 mins, the cells were centrifuged at 1000xG 2 mins, washed in cold PBS before repelletting and lysis in 60µl HNTE.

As control samples, the remaining un-electroporated cells in the pS65Ub and HA-Ub mixtures were transferred to separate 500µl DMEM aliquots (sample ‘15 - electroporation’). Additionally, two smaller cell aliquots (2x 6.25x10^5^) were resuspended in 62.5µl Buffer R alone. For one suspension, 6x10µl was electroporated then combined in a 500µl DMEM aliquot (sample ‘0 + electroporation’), for the other, 60µl was transferred directly into a DMEM aliquot (sample ‘0 - electroporation’). All control samples were harvested after 15 mins incubation at 37°C.

### Affinity purification mass spectrometry (APMS) sample preparation

Eluted pS65Ub APMS and nucleosome APMS samples were reduced (4.5mM DTT, 20 mins at 60°C), alkylated (11mM IAA, 20 mins at RT in the dark) then quenched (16.5mM DTT). Protease digestion was performed with incubation with 300ng trypsin at 37°C overnight, or incubation with 100ng Lys-C at 37°C for 1.5 hrs followed by addition of 100ng trypsin and overnight incubation. The digested samples were acidified with 20% trifluoroacetic acid (TFA), desalted using C18 stage-tips as previously described^129^, then dried. The samples were then labelled with TMT11-plex (pS65Ub APMS) or TMTpro 16-plex (nucleosome APMS) reagents. TMT labels were dissolved in anhydrous acetonitrile and peptide samples reconstituted pH8.5 HEPES and mixed to a final concentration of 40% ACN (TMT11-plex) or 28% ACN (TMTpro 16-plex), incubated at room temperature for 1 hr then quenched by addition of hydroxylamine (5%v/v, 15 mins RT). TMT-labelled peptides were then combined 1:1 and dried down.

The pS65Ub APMS samples were resuspended in 0.1% TFA, desalted as before, then dissolved in Buffer A (2% acetonitrile, 0.1% formic acid) before mass spectrometry analysis. The nucleosome APMS samples were subjected to reverse phase separation using stage tips packed with Empore SPE Disks as per manufacturers instructions with the following modifications: the sample was resuspended and loaded onto the column in 2% acetonitrile 5% formic acid then washed twice in 100 uL 2% acetonitrile 1% formic acid and three times in 1% formic acid. Four fractions were eluted using the following buffers: 7.5% ACN/25 mM NH4COOH pH10; 12.5% ACN/25 mM NH4COOH, pH 10; 17.5% ACN/25 mM NH4COOH, pH10; 80% ACN/2% NH4OH, pH10. The eluates were then dried down and resuspended in Buffer A for mass spectrometry analysis.

### Affinity Purification Mass Spectrometry Analysis

LC-MS/MS analysis was performed on an Orbitrap Eclipse mass spectrometer (Thermo) coupled to a Dionex Ultimate 3000 RSLC (Thermo) employing a 25 cm IonOpticks Aurora Series column (IonOpticks) with a gradient of 2% to 30% Buffer B (98% ACN, 2% H2O with 0.1% FA, flow rate = 300 nL/min) in Buffer A.

For pS65Ub APMS, the total run time was 185 mins. The Orbitrap Eclipse collected FTMS1 scans at 120,000 resolution with a 50 ms maximum injection time and 250% normalized AGC target. FTMS2 scans on precursors with charge state of 2-6 were collected at 15,000 resolution with CID fragmentation at a normalized collision energy of 35%, normalized AGC target of 150%, and max injection time of 100 ms. For nucleosome APMS, the total run time was 65 mins . The Orbitrap Eclipse with FAIMS Pro DUO of -50, -70CV collected FTMS1 scans at 240,000 resolution with a 50 ms maximum injection time and 250% normalized AGC target. FTMS2 scans on precursors with charge states of 2-4 were collected at 15,000 resolution with HCD fragmentation at a normalized collision energy of 30%, normalized AGC target of 300%, and max injection time of 11 ms.

For nucleosome APMS fractions, real-time database search (RTS) was performed prior to acquisition of MS3 spectra using InSeqAPI software^130^, operating as published previously^131,132^. The RTS parameters follow: Uniprot human database August 2021 version (with common contaminants and decoys, and 218,136 Swissprot sequences of canonical and protein isoforms); the static modifications were K and n-term TMT (+229.162932), C carbamidomethylation (+57.0215); the variable modifications were M oxidation (+15.9949) and Y TMT (+229.162932). Protein closeout was employed, requiring 3 distinct and 5 unique peptides across runs. Comet v.2019.01 was used for offline searching with the same parameters. The pS65Ub APMS samples were searched offline only against Uniprot human database April 2021 version, otherwise using the same parameters.

We used a linear discriminator algorithm to filter peptide FDR to <1%, and an in-house software package to quantify the highest peak within 20 ppm of theoretical TMT reporter ion mass windows and correct for isotope impurity. Statistical testing and quantification were performed by MSstatsTMT_2.0.1 R package^133^. Median normalization across channels facilitated systematic bias correction. For nucleosome APMS, multiple fractions were combined in the software, and where a species was found in more than 1 fraction, the fraction with the highest maximal reporter ion intensity was utilized.

### Targeted Mass Spectrometry

Purified histone samples were prepared as described in the ‘Histone Purification’ section. 1ug of each histone prep sample was subjected to overnight enzymatic digestion at 37°C with sequencing grade trypsin (Promega, enzyme : protein ratio = 1:50). Resultant peptides were acidified with 20% Trifluoroacetic acid (TFA, 1% final concentration), and desalted using C18 stage-tips^129^. A mixture of ubiquitin AQUA peptides were spiked into each sample at 1pmol per peptide and the samples were lyophilized, and re-suspended in 10 μL Buffer A (2% ACN, 0.1% formic acid) for targeted LC-MS/MS analysis.

Targeted LC-MS/MS analysis was performed by injecting 1μL of each sample on an Orbitrap Ascend mass spectrometer (ThermoFisher) coupled to a Dionex Ultimate 3000 RSLC (ThermoFisher) employing a 25 cm IonOpticks Aurora Series column (IonOpticks, Parkville, Australia) with a gradient of 2% to 30% buffer B (98% ACN, 2% H2O with 0.1% FA, flow rate = 300 nL/min). The targeted samples were analyzed with a total run time of 95 min. For all samples, the Orbitrap Ascend collected FTMS1 scans at 120,000 resolution with an AGC target of 1 x 10^6^ and a maximum injection time of 50 ms. FTMS2 scans on precursors with charge states of 3-6 were collected at 15,000 resolution with CID fragmentation at a normalized collision energy of 30%, an AGC target of 2 x 10^4^, and max injection time of 400 ms.

### ChIP Seq

ChIP Seq was performed as previously described^134^ with the following modifications: 50mM DSG (disuccinimidyl glutarate, #C1104 - ProteoChem) for 30 mins followed by 10 mins of 1% formaldehyde was used to double crosslink HeLa or iPSC-derived NGN2 neurons (5x10^6^), followed by the addition of glycine to quench formaldehyde. Nuclei were released using ChIP Nuclear Isolation Buffer (20 mM Tris pH 8.0, 1% Triton X-100, 0.1% SDS, 150 mM NaCl and 1mM EDTA), and chromatin sheared via sonication (Covaris sonicator: Fill level – 10, Duty Cycle – 15, PIP – 350, Cycles/Burst – 200, Time – 8-mins).

Solutions containing sheared chromatin were mixed with relevant antibodies and rotated at 4°C overnight before protein A magnetic beads were then added to isolate antibody-bound chromatin. The beads were washed twice in each of the following buffers: ChIP Wash Buffer #1 (20 mM Tris pH 8.0, 1% Triton X-100, 0.1% SDS, 150 mM NaCl, 1 mM EDTA, and 0.1% NaDOC), ChIP Wash Buffer #2. (20 mM Tris pH 8.0, 1% Triton X-100, 0.1% SDS, 500 mM NaCl, 1 mM EDTA, and 0.1% NaDOC) and ChIP Wash Buffer #3 (0.25 M LiCl, 0.5% NP-40, 1mM EDTA, 20 mM Tris, pH 8.0, 0.5% NaDOC), followed by single wash in TE buffer (200 mM Tris pH 8.0 and 10 mM EDTA). The mixtures were then shaken for 20 mins in Elution Buffer (100mM NaHCO_3_, 1% SDS), before overnight incubation at 65°C to de-crosslink DNA. The DNA fragments were purified with MinElute PCR purification kit (Qiagen), then quantified by Qubit. Libraries were constructed with the Ovation Ultralow Library System V2 (Tecan) using 2ng of each sample according to the manufacturer’s instructions, then sequenced using paired-end 50bp read configuration.

The ENCODE analysis pipeline (v2.2.1)^135^ was used, with Bowtie2 (v2.3.4.3)^136^ employed to align reads to the reference genome (hg38). Duplicated or low-quality reads were filtered out using Picard (Broad Institute - v2.20.7) and samtools (v1.9)^137^, with further filtering to remove regions where genome assembly introduces erroneous signal^138^. The SPP data processing pipeline was used to call peaks, with the level of background signal assessed via reference to input sample^139^. DiffBind (v3.12.0) was used to identify differential peaks as follows. ChIP Seq bam and bed files were loaded with the dba() function to create Differential Binding Analysis object^140^, followed by read counting with the dba.count() function then depth normalization using the dba.normalize() function. The dba.analyze() function was used to call differential peaks across the conditions using DESeq2 (FDR<0.01). ChIPseeker (v1.38.0) was utilized to identify nearest genes^141^. RPKM-normalized ChIP Seq data was visualized using IGV (2.18.4) genome browser.

### CUT&RUN

CUT&RUN was performed using the CUTANA ChIC/CUT&RUN Kit (Epicypher) as per manufacturers instructions with the addition of DUB inhibitor PR-619 (100µM) to Wash and Antibody buffers. Indexed libraries generated using NEBNext library prep kit (NEB) and NEBNext Multiplex Oligos for Illumina (NEB) as per manufacturers instructions. Paired-end reads were aligned to the reference genome (hg38) using Bowtie2 (-p 24 -X2000 -t -x) and sorted with Samtools (view -bS -F 4 -q 30 -u - | sort -T). PCR duplicates were marked using Picard MarkDuplicates (MAX_FILE_HANDLES_FOR_READ_ENDS_MAP=1000 REMOVE_DUPLICATES=true ASSUME_SORTED=true VALIDATION_STRINGENCY=LENIENT). Genome-wide coverage was determined using Bedtools genomecov (-ibam -bg -scale -g), converted to a BigWig file with wigToBigWig -clip, and MACS2 (callpeak -t --outdir . -n -f BAM -g hs -p 0.00001 --nomodel --keep-dup all) was used for peak calling.

The metagene profiles were composed of the following: (blue) 948 pS65Ub-positive genes with annotation in GENCODE v32 and (red) 1,000 randomly selected genes as a negative control.

### Bulk RNA Seq Analysis

RNA was purified from ∼5x10^6^ WT and PINK1 knockout HeLa cells per sample using the RNeasy Plus Mini Kit (Qiagen) as per manufacturers instructions. RNA Seq library generation and sample analysis was performed as described previously^142^ with the following modification: prior to alignment to the human reference genome, the lowest expressing 70% of the features were excluded.

To identify candidate genes showing pS65Ub-dependent expression in HeLa cells, we isolated genes with reduced OA-dependent expression upon PINK1 KO. To do this, we subtracted WT from KO log_2_(OA/V) fold change values. WTlog_2_(OA/V) - KOlog_2_(OA/V) = ΔFC. Genes with a ΔFC value >0.5 were considered to be unregulated in WT cells.

For analysis of dopaminergic neuron samples (see ‘Dopaminergic Neuron Transduction’ for sample generation), RNA was purified from 1x10^6^ cells per sample, then library generation and sample analysis was performed as described above. Three independent replicates were conducted for each transduction condition (iPink1 WT, iPink1 KD, and BFP). One BFP replicate was removed from the analysis after PCA identified it as a clear outlier, suggesting technical variability rather than a biological effect.

To identify overrepresented gene sets in WT, KD or BFP-transduced dopaminergic neurons, we isolated significantly up and down-regulated genes (FDR <0.5 and LogFC >0.5 or <-0.5 respectively) from Extended Data 13-15 then performed an over-representation analysis using the hypergeometric test, using the term set MSigDB Collection C2 Subset CGP and a minimum set size of 15. Only genes upregulated in WT vs KD conditions (Extended Data 13) returned significantly enriched genesets.

### ATAC Seq Analysis

Wild type HeLa cells were treated with OA (1µM, 24 hrs), then bulk ATAC Seq according to manufacturers instructions (Active Motif, #53150). 50,000 cells per condition were used to generate libraries.

### Library Sequencing

Libraries were sequencing using the Illumina sequencing platform (NovaSeq 6000, NovaSeq XPlus, NextSeq 2000).

### GSEA Analysis

To analyze pS65Ub APMS data, we used the R package ‘fgsea’, with minimum set size = 10. For CUT&RUN and ChIP Seq data, called peaks were annotated using ChIPseeker (tssRegion=c(-1000, 1000)). The set of genes with peaks annotated as promoter-proximal were used for GSEA. The enrichr module of GSEApy was used to determine genset enrichment against the MSigDB Human Gene Ontology (GO) Biological Process (BP) collection (2023).

### Seahorse XF Mito Stress Test

HeLa cells doxycycline-inducibly expressing iPink1 WT or KD were plated on day 1 on Seahorse XF Pro M Cell Culture Microplates (#103774-100) in basal growth media described above. Where indicated, cells were plated in media containing 1µM (0.45µg/ml) doxycycline. On day 2, the cells were switched to no glucose DMEM supplemented with 10% FBS, 4 mM glutamine, 1mM pyruvate and either 25mM glucose or 10mM galactose, supplemented with 1µM doxycycline where indicated. Cells were grown in glucose- or galactose-containing medium (+/- doxycycline) for 72 hours before analysis.

Seahorse XF Mito Stress Test Kit (Agilent Technologies #103015-100) was used according to the manufacturer’s protocol and run with the Seahorse XFe96 Analyzer (Agilent Technologies). One hour prior experiment, cells were switched to Seahorse XF DMEM Medium, pH 7.4 (Agilent technologies #103575-100) supplemented with 10% FBS, 4 mM glutamine, 1mM pyruvate and either 25mM glucose or 10mM galactose incubated 37C in the absence of CO_2_ . Compounds were used at the following concentrations: Oligomycin 1.5uM, FCCP 2.0µM, rotenone/antimycin A 0.5µM. Each experiment is normalized by the number of cells as counted per Hoechst 33342 staining using the Agilent BioTek Cytation 1 (Agilent Technologies). Analysis was performed with the Seahorse Wave Controller Software 2.6 (Agilent Technologies) exporting the data with the report generator provided with the software.

### H2AK119ub antibodies epitope testing

To confirm that H2AK119ub antibody binding was unaffected by phosphorylation of ubiquitin at serine 65, we phosphorylated H2AK119ub nucleosomes with constitutively active *Tribolium castaneum* Pink1 (insect Pink1/iPink1) *in vitro* then assessed H2AK119ub antibody recognition by Western blot. First HEK293T were transfected with TET ON 3xNES-iPink1-V5 WT or KD, then incubated with 1µg/ml doxycycline to drive iPink1 expression. After 16 hours incubation, cells were lysed in HNTE, centrifuged at 21,130xG (5 mins, 4°C), then post-nuclear supernatants were rotated with 50µl (1mg) V5-Trap Magnetic Agarose at 4°C for 4 hrs, washed with 3x1ml 300mM NaCl HNT, then resuspended in 100µl Kinase Reaction Buffer (KRB: 20 mM HEPES pH 7.5, 20 mM MgCl2, 2 mM dithiothreitol, 1X HALT). To perform the first set of kinase reactions, two 4µl aliquots of stably pre-ubiquitnated nucleosomes were diluted into 46µl V5 bead-bound WT and KD iPink1 mixtures supplemented with 2mM ATP and incubated at 30C for 30 minutes.

We performed a second set of kinase reactions in parallel on nucleosomes ubiquitinated in house. During the 4 hrs incubation at 4°C, H2AK119ub nucs were generated as described in the ‘*In Vitro* Ubiquitination and Deubiquitination Assays’ section, using wild type FLAG ubiquitin as the Ub source, 16µg non biotinylated nucleosomes, 3µM RING1B-PCGF1, 200µl reaction mixtures and incubating the full ligase reaction at room temperature for 40 mins. The ligase reaction mixtures were then split into two equally sized aliquots. The remaining V5 bead-bound WT and KD iPink1 mixtures were immobilised on a magnet, and the ligase reaction mixtures were used to resuspend the beads. The mixtures were moved to 30C to begin the kinase reaction, and were gently resuspended once at 20 minutes due to bead settling.

Stably pre-ubiquitinated (set 1) and in house ubiquitinated (set 2) reactions were killed with addition of LDS sample buffer after 30 mins to allow sample analysis by immunoblot.

### Dopaminergic Neuron Transduction

We designed lentiviruses encoding wild type or kinase dead 3xNES-iPink1-V5, or blue fluorescent protein (BFP), all under an EF1a promoter and encoding a EGFP marker connected by a T2A tag to track transduction rates. Lentiviral particles were generated by Vectorbuilder. We then transduced immature dopaminergic neurons at MOI=5 on differentiation day 22, performing a full media change 24 hours later.

For immunofluorescence experiments shown in Figure 7A, dopaminergic neurons were transduced on differentiation day 22 or treated with OA or vehicle for the final 24 hrs before fixation at day 53. In Extended Data Figure 7D/E, dopaminergic neurons transduced on differentiation day 29 before analysis on day 53. Bulk RNA Seq, scRNA Seq and ChIP Seq experiments were performed in parallel on cells transduced at differentiation day 22 and harvested at day 62.

### Single Cell Dissociation of Mature Dopaminergic Neurons

Mature cultures of dopaminergic neurons were first washed with 37°C PBS before being treated with 37°C dissociation media containing Accutase and Papain dissolved in PBS at 20 Units / mL, in a 1:1 ratio, for 5 mins at 22°C. Dissociation Buffer was then quenched with an equal volume of warm culture media containing 10µM Y27632 and a 0.15 Units / mL solution of DNase1. The cells were then pooled and triturated using a 1 mL pipette with standard bore tips to dissociate the network of neurites from the somas. The dissociated mixture was then passed through a 40µM cell strainer and centrifuged at 150g for 3 mins. The supernatant was then aspirated and the pellet was gently resuspended in room temperature Wash Buffer composed of PBS and 0.2% BSA before centrifuging again at 150g for 3 mins. This wash step was repeated once more before passing the final sample through a 40µm strainer. Samples were kept on ice as preparations were made for loading.

### scRNA Seq Processing

Single cell suspensions loaded in a concentration of 1500 cells/µl on a 10x Genomics chip, and then run on a Chromium X Controller which encapsulated single cells into gel beads in emulsion (GEMs). Emulsions were processed into amplified cDNA products following user guide Chromium GEM-X Single Cell 3’ Reagent Kits v4 CG000731.

Library construction was performed according to user guide specifications with 10 cycles of amplification for SIPCR. For each final library, average size and concentration were evaluated using Agilent Technologies TapeStation 4200 and ThermoFisher Scientific Qubit Fluorometer respectively. The final DNA libraries were paired end sequencing on a Novaseq X+ from Illumina Technologies (San Diego, CA) using a 10B 100 cycle kit with the following sequencing specifications (28 cycles: Read 1, 10 cycles: i7 index, 10 cycles: i5 index and 90 cycles: Read 2) with a targeted sequencing depth of 20,000, read pairs per cell. Raw sequencing data was demultiplexed by Illumina’s Bcl2FASTQ software then resultant FASTQ data was run through cellranger-9.0.0 software to generate gene expression matrix for downstream analysis.

### scRNA Seq Analysis

Sequencing data were analyzed using the Cell Ranger software suite (version 7.1.0). The workflow included the following steps: Raw FASTQ reads were aligned to the custom human reference genome GRCh38 with EGFP. The Gene expression was quantified using Cell Ranger’s default settings, excluding introns. For downstream processing of the count data with Seurat(V5.0.1)^143^ using the default settings of the package. For cell level QC, cells with <500 unique detected genes, >10% mitochondrial UMIs and < 1 UMI count of eGFP were discarded. After the filtering step, the gene × cell matrix of raw UMI counts was log-normalized using ‘NormalizeData()’ and scaled using ‘ScaleData() ’using Seurat(V5.0.1). Then, performed dimensionality reduction by PCA, calculated UMAP coordinates and Louvain clustering for all cells using Seurat(V5.0.1). The final clusters were annotated using markers described from Jerber et al. 2021 and Bressan et al. 2023^95,144^. For calculating gene set scores for single cell datasets, raw counts were converted to counts-per-million with edgeR^145^, then transformed as log_2_(CPM+1). For each cell, we summarized the set by the arithmetic mean of log_2_(CPM+1) across genes present in the set, yielding one continuous score per sample and gene set. For testing whether a gene set is enriched in a cell type, we performed GSEA and reported normalized enriched score, nominal p value and adjusted p value^146,147^.

## Acknowledgements

We thank Carlos Sanchez Priego for his help in generating iPSC-derived neurons, and both Andrea Cochran and Asad Taherbhoy for providing RING1B-PCGF proteins for use in in vitro ubiquitination assays. We thank Ai Zhang for her insights on our neuronal image analysis, and Meng He for providing engineered cell lines. Thanks also to Andrea Cochran, Samuel Killackey, Alessandro Ori and Antonina Hafner for their helpful feedback while preparing this manuscript. We thank the Neuropathology Laboratory and Brain Bank at Mayo Clinic Jacksonville for processing human post-mortem tissues and the excellent technical support. We are grateful to the patients and their families who made this study possible. Research in the Springer lab is supported by the National Institutes of Health (NIH)/National Institute of Neurological Disorders and Stroke (NINDS) [RF1NS085070, R01NS110085, and U54NS110435], the Department of Defense Congressionally Directed Medical Research Programs (CDMRP) [W81XWH-17-1-0248], The Michael J Fox Foundation for Parkinson’s Research, The Ted Nash Long Life Foundation, Mayo Clinic Foundation, and the Mayo Clinic Robert and Arlene Kogod Center on Aging (all to W.S.). F.C.F. is supported by the Research Catalyst Award from the Center of Biomedical Discovery and the Office of Belonging at the Mayo Clinic. Mayo Clinic Jacksonville is an American Parkinson Disease Association (APDA) Center for Advanced Research (D.W.D. and W.S.).

## Author Contributions

T.J.M. conceived of the study, designed and performed the experiments, analysed the data and authored the manuscript. B.D. performed (with the help of V.K.), analyzed and helped design ChIP seq experiments and contributed to experimental strategy and manuscript authoring. C.F. cultured, treated and harvested iPSC-derived neurons and C.F. and C.J. helped design relevant experiments. D.L. analysed RNA seq and CUT&RUN datasets, S.L., H.N. and O.F. optimized, performed and analysed the pS65Ub immunohistochemistry analysis. V.V.K. captured immunofluorescence images of iDopa samples and performed live cell imaging. X.H., F.C.F and W.S. designed, performed and analyzed fibroblast experiments and PD tissues super resolution microscopy analyses. M.J. analyzed scRNA Seq data, and T.K.C. performed targeted mass spectrometry analyses. B.J.R., R.H.R. and A.J.S. performed α-synuclein seeding experiments as well as immunofluorescence analyses of PINK1/Parkin mutant neurons. D.W.D identified PD tissues for super resolution microscopy analysis. D.G. helped with Seahorse assays. C.M.R. and B.B. supervised the work and contributed to both experimental design and manuscript authoring. Research in the Ryan Lab is supported by the Medical Research Council grant (MR/Y014987/1), Rosetrees Trust (PhD2024\100031), Lady Margaret Hall Alison Brading Scholarship and The Clarendon Fund.

---

**Extended Data Figure 1.**
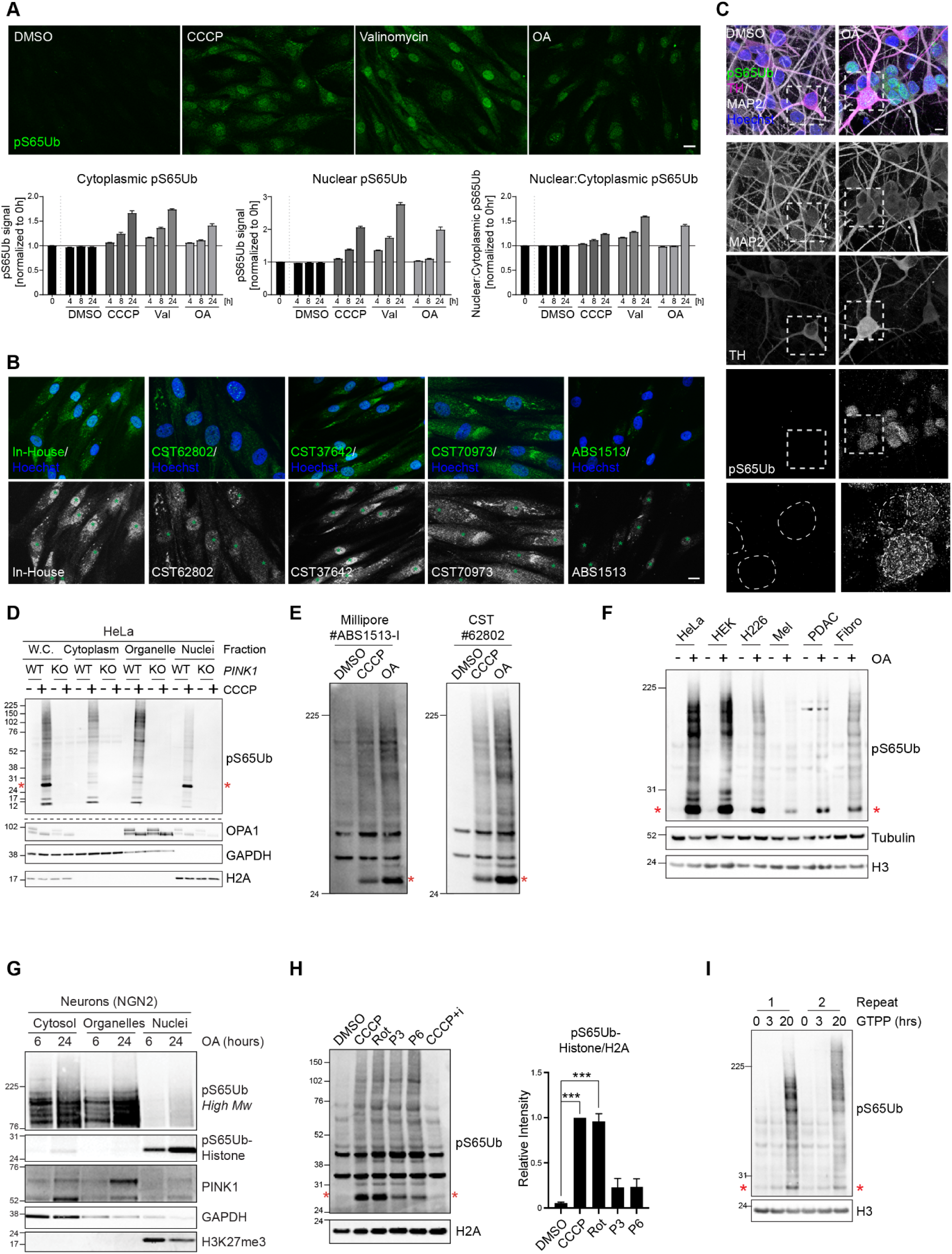
Both nuclear pS65Ub and pS65Ub-histones are widely detectable. **A** - Human fibroblasts were treated with mitochondrial toxins (CCCP, 50µM; valinomycin, 1µM; oligomycin 10µM + antimycin, 4µM) or DMSO for 0, 4, 8 and 24 hrs. Top: Representative immunofluorescence images from 24 hr timepoint showing distribution of pS65Ub (green). Bottom: Quantification shows mean +/- SEM pS65Ub intensity in cytoplasm, nucleus and nucleus:cytoplasm relative to 0 hrs timepoint (separated with vertical dashed line) from 1 experiment with 3 technical repeats. The horizontal line shows the 0hr-normalized baseline. **B** - Fibroblasts were treated with valinomycin (1µM, 24 hrs) before pS65Ub (green) was visualized by immunofluorescence using multiple specific antibodies. Nuclei (blue) are annotated with green asterisks. **C** - Confocal Z-stack max projections showing human induced dopaminergic neurons treated with oligomycin + antimycin (OA, 1µM, 24 hrs) and stained for pS65Ub (green), the dopaminergic neuron marker tyrosine hydroxylase (TH; magneta), MAP2 (white) and Hoechst (blue). Dashed boxes indicate areas magnified in pS65Ub channel inset and dashed circles delineate nuclei. **D** - Wild type (WT) and PINK1 knockout (KO) HeLa cells were treated with CCCP (20µM, 4 hrs) as indicated, then subcellular fractionations were analysed by Western blot. **E** - HeLa were treated with CCCP (20µM, 4 hrs) or OA (1µM, 24 hrs) before immunoblot analysis of whole-cell lysates using two pS65Ub antibodies. **F** - HeLa, HEK293T, NCI-H226, murine melanoma, murine PDAC or human skin fibroblast cells were treated with OA, then pS65Ub levels were assessed by Western blot. **G** - NGN2-induced iPSC-derived neurons were treated with OA (1µM, 6 or 24 hrs) before subcellular fractionation and Western blot analysis. **H** - HeLa were treated with CCCP (20µM, 4 hrs), rotenone (Rot - 20µM, 20 hrs), paraquat (P3/P6 - 3/6mM, 20 hrs) or CCCP+i (PINK1 inhibitor/PRT062607, 2.5µM, 4 hrs). Quantifications show mean +/-SEM from three independent repeats, *** p<0.0001 (CCCP+i treatment performed twice only so excluded from quantification). **I** - HeLa cells were treated with 10µM Gamitrinib-TPP (GTPP) for 0, 3 or 20hrs before analysis of whole-cell lysates by Western blot. Red asterisks mark the pS65Ub-histone band. Scale bars 10µm (50µM for panel D). Dashed lines in panels E and F indicate where identical samples were run on separate blots.

**Extended Data Figure 2.**
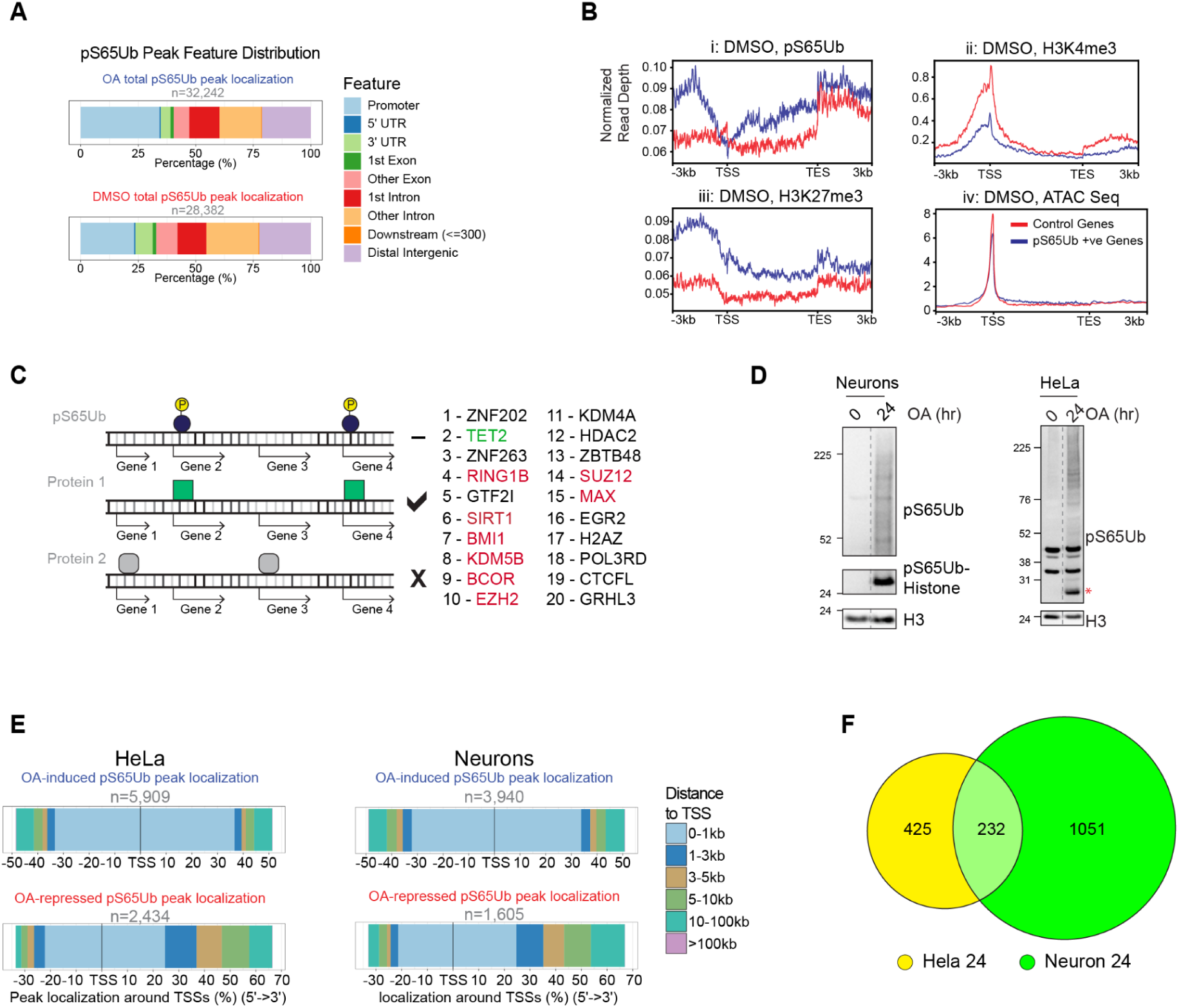
Characterization of pS65Ub-positive chromatin domains. **A** - Representative genomic feature enrichment of pS65Ub CUT&RUN peaks from DMSO and OA (1µM, 24 hrs) conditions. **B** - HeLa cells were treated with OA or DMSO before CUT&RUN and ATAC Seq analysis. Averaged pS65Ub (i), H3K4me3 (ii) and H3K27me3 (iii) CUT&RUN signal, or ATAC Seq signal (iv) from the DMSO condition are displayed as metagene plots. pS65Ub-positive (blue, n=948) and control gene sets (red, n=1000) genes aligned with gene transcription start (TSS) and end (TES) sites, are displayed. Experiments i, ii/iii and iv were performed independently. **C** - Schematic description of coregulator analysis. Proteins that bind similar genes to pS65Ub were identified (**Extended Data Table 5**). The top 20 are listed with TET2 highlighted in green and polycomb-associated proteins in red. **D** - Input samples for pS65Ub ChIP experiments in neurons and HeLa. Cells were treated with OA for 0 or 24 hrs as indicated before Western blot analysis of whole-cell lysates. A red asterisk marks the pS65Ub band for HeLa samples. **E** - Quantification of distance to nearest TSS of pS65Ub ChIP Seq peaks showing induction vs repression after OA treatment in HeLa cells (left) and neurons (right). **F** - Venn diagram showing overlap between genes displaying OA-dependent pS65Ub enrichment at the TSS in both HeLa cells and neurons.

**Extended Data Figure 3.**
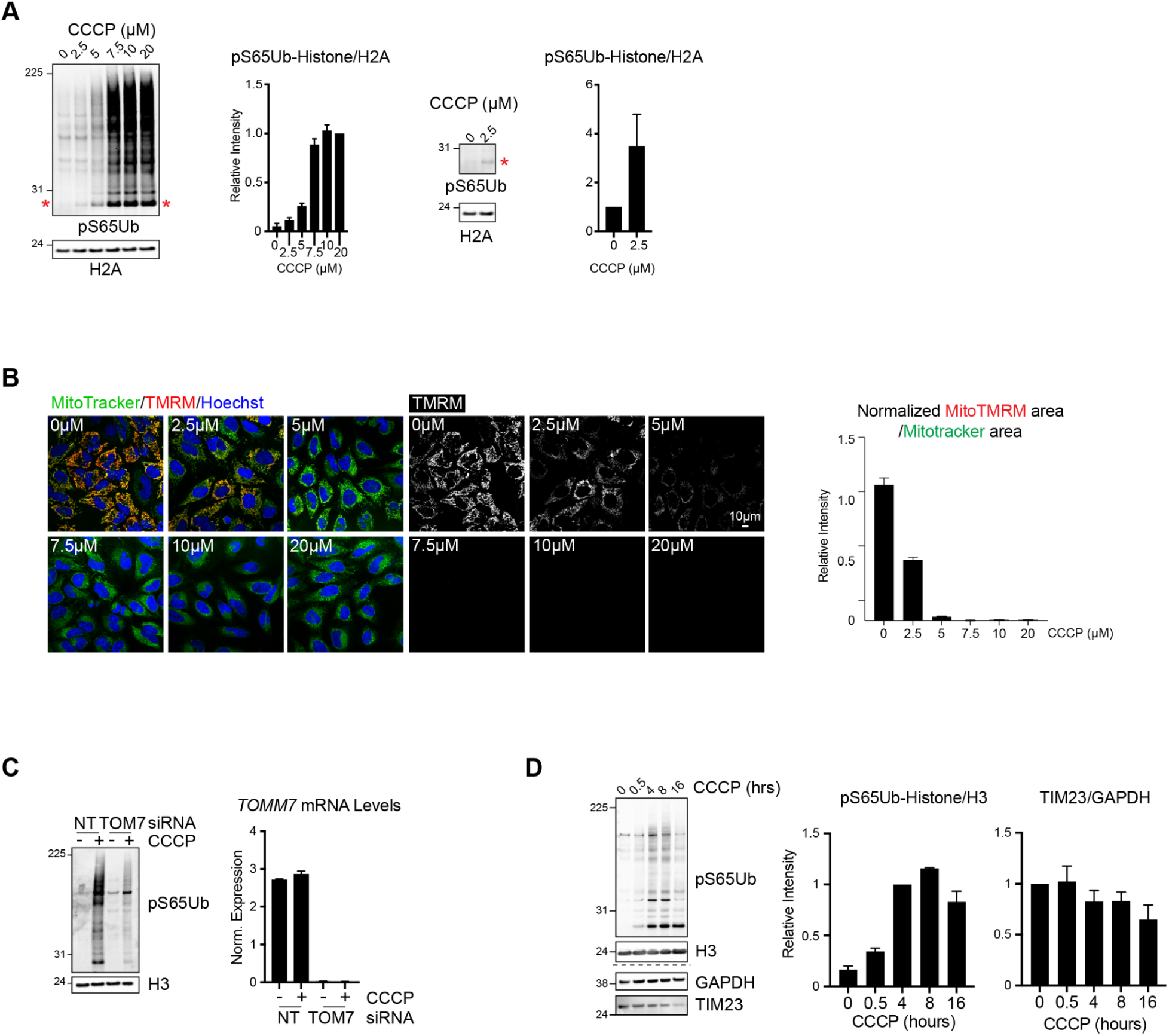
Histone pS65 ubiquitination corresponds with PINK1 mitochondrial recruitment. **A** - HeLa were treated with a dose range of CCCP for 4 hrs. Histone pS65Ubiquitination was quantified from three independent repeats (mean +/- SEM). The 0µM and 2.5µM CCCP conditions are presented alone (right) to demonstrate the degree of histone pS65Ubiquitination at ‘sub-threshold’ CCCP concentrations. Red asterisks mark the histone-pS65Ub band. **B** - HeLa treated as above were stained with MitoTracker (green) or MitoProbe TMRM (red before live cell imaging. TMRM/MitoTracker ratios were calculated to measure relative mitochondrial polarization. Mean +/-SEM from 3 independent repeats. **C** - HeLa cells transfected with TOM7-specific or non-targeting (NT) siRNAs then treated with CCCP (4 hrs, 20µM). Loss of TOM7-dependent PINK1 activation was confirmed by Western blot (left) and *TOMM7* mRNA depletion was demonstrated by qPCR (right). **D** - HEK293T cells were treated with CCCP (20µM) for 0-16 hrs as indicated before analysis of whole-cell lysates by Western blot. Normalized pS65Ub-Histone and TIM23 levels were quantified from three independent repeats (mean +/- SEM).

**Extended Data Figure 4.**
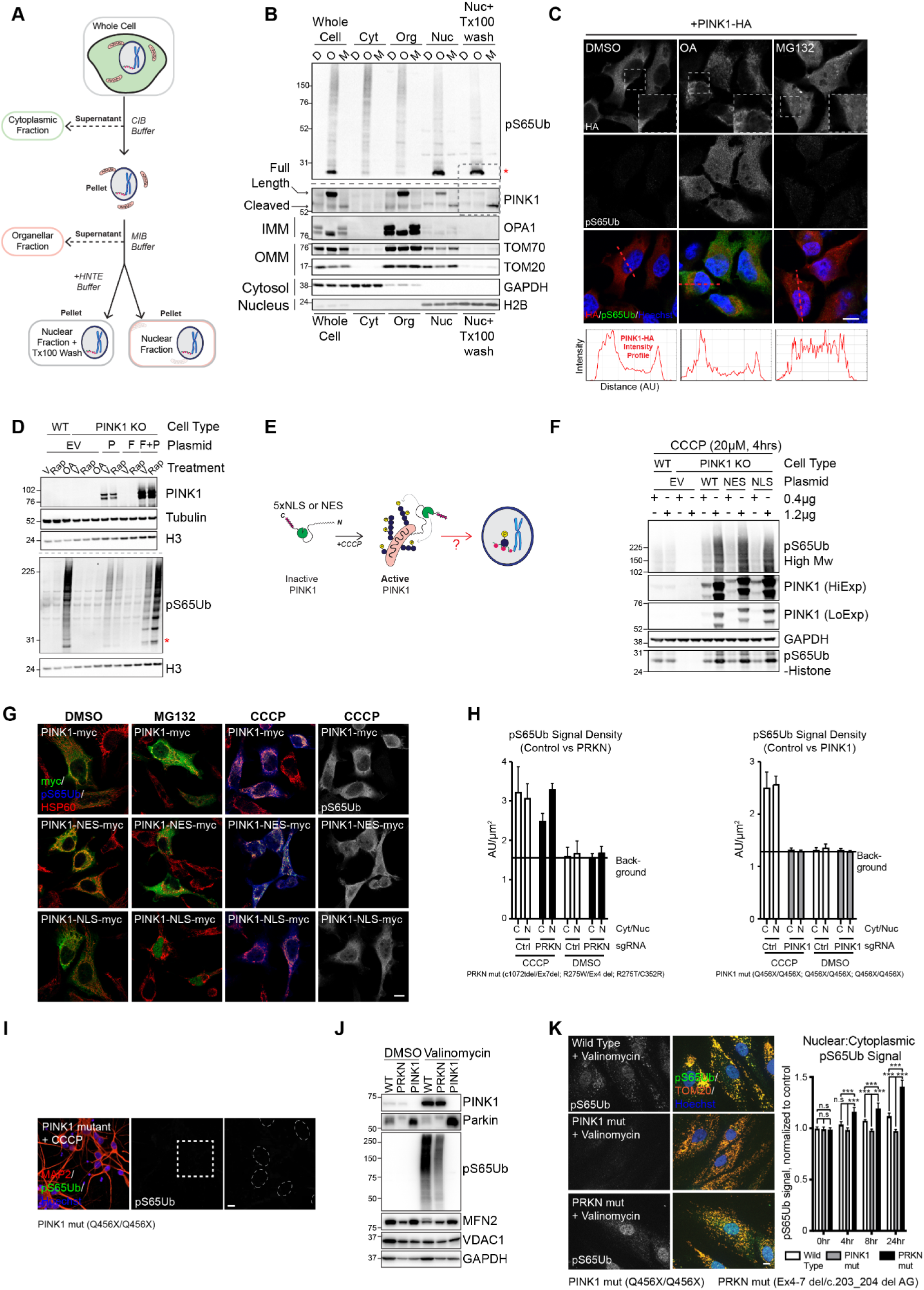
The nuclear pS65Ub pool is regulated by the activities of PINK1 and Parkin at mitochondria. **A** - Schematic depiction of subcellular fraction experiment performed in Extended Data Figure 3B. Detergent buffers used for fractionation are given in italics. CIB and MIB buffers are from Cell Fractionation Kit (Cell Signaling Technologies, #9038), while HNTE was made in house. **B** - Cells were treated with OA (1µM, 18 hrs), MG132 (20µM, 4 hrs) or DMSO before subcellular fractionation and Western blotting. Protein subcellular localizations are annotated (IMM = inner mitochondrial membrane, OMM = outer mitochondrial membrane) and arrows indicate both full length and cleaved PINK1 bands. For the pS65Ub blot, identical samples were analyzed on a separate gel (separated by dashed lines). **C** - PINK1-HA was transiently expressed in HeLa cells before treatment with OA, MG132 or DMSO as before. The distributions of pS65Ub (green), HA (red) and DNA (blue) were assessed by immunofluorescence. Magnified regions of interest are indicated by dashed boxes. Red dashed lines indicate regions captured by intensity profile (performed in the red/HA channel), graphed below. **D** - WT or PINK1 KO HeLa were transiently transfected with empty vector (EV), PINK1-FKBP (P) or FIS1-FRB (F) in combinations indicated before 18 hrs treatment with OA (1μM) or rapalog (500nM). A dashed line reveals where identical samples were analyzed on a separate gel. **E** - Schematic depiction of experiments performed in Extended Data Figures 4F, G. **F** - WT or PINK1 KO HeLa cells were transiently transfected with EV, PINK1 WT, PINK1-NES or PINK1-NLS before treatment with CCCP and Western blot analysis. Two plasmid amounts were used for transfection to achieve high and low relative PINK1 expression, captured by high and low exposures (HiExp/LoExp respectively). **G** - PINK1-myc tagged with NES/NLS signals were transiently expressed in HeLa cells before treatment with CCCP (20µM, 4 hrs) or MG132 (20µM, 4 hrs). Cells were fixed and stained for myc (green), pS65Ub (blue, grayscale in the far right column) or mitochondrial marker HSP60 (red). Nuclear exclusion (NES) and sequestration (NLS) is observed after MG132 treatment. **H -** Quantification of nuclear vs cytoplasmic pS65Ub signal density (arbitrary units/µm^2^) in healthy control, *PRKN* (left) and *PINK1* (right) mutant iPSC-derived dopaminergic neurons after treatment with CCCP (10µM, 6 hrs). The horizontal line shows the signal on DMSO-treatment, considered background due to negligible PINK1 activation/pS65Ub levels. Mean +/- SEM from three independent cell lines per genotype. **I** - iPSC-derived dopaminergic neurons from PD patients with mutations in *PINK1* were treated with CCCP (10µM, 6 hrs). Dashed boxes annotate the area magnified in the inset and dashed circles show nuclear borders. Inset zoom in of pS65Ub channel. See Figure 3G for Control and *PRKN* mutant conditions. **J** - Skin fibroblasts from *PINK1*/*PRKN* mutant PD donors and healthy controls were treated with valinomycin (1µM, 8 hours) before Western blot analysis. Maximal Parkin activity (revealed by MFN2 ubiquitination, see band shift) and PINK1 activity (substrate pS65Ubiquitination) is only detected in WT cells treated with valinomycin. **K** - Fibroblasts were treated with valinomycin (1µM, 0-24 hrs) before fixation and immunofluorescence analysis of pS65Ub (green, grayscale in the left hand column), TOM20 (orange) and DNA (blue). Nuclear:cytoplasmic pS65Ub signal intensity from three experiments was quantified, with mean +/-SEM, 2-way ANOVA shown. Red asterisks identify the pS65Ub-histone band *p<0.05, **p<0.01, ***p<0.001. Scale bars 10µm.

**Extended Data Figure 5.**
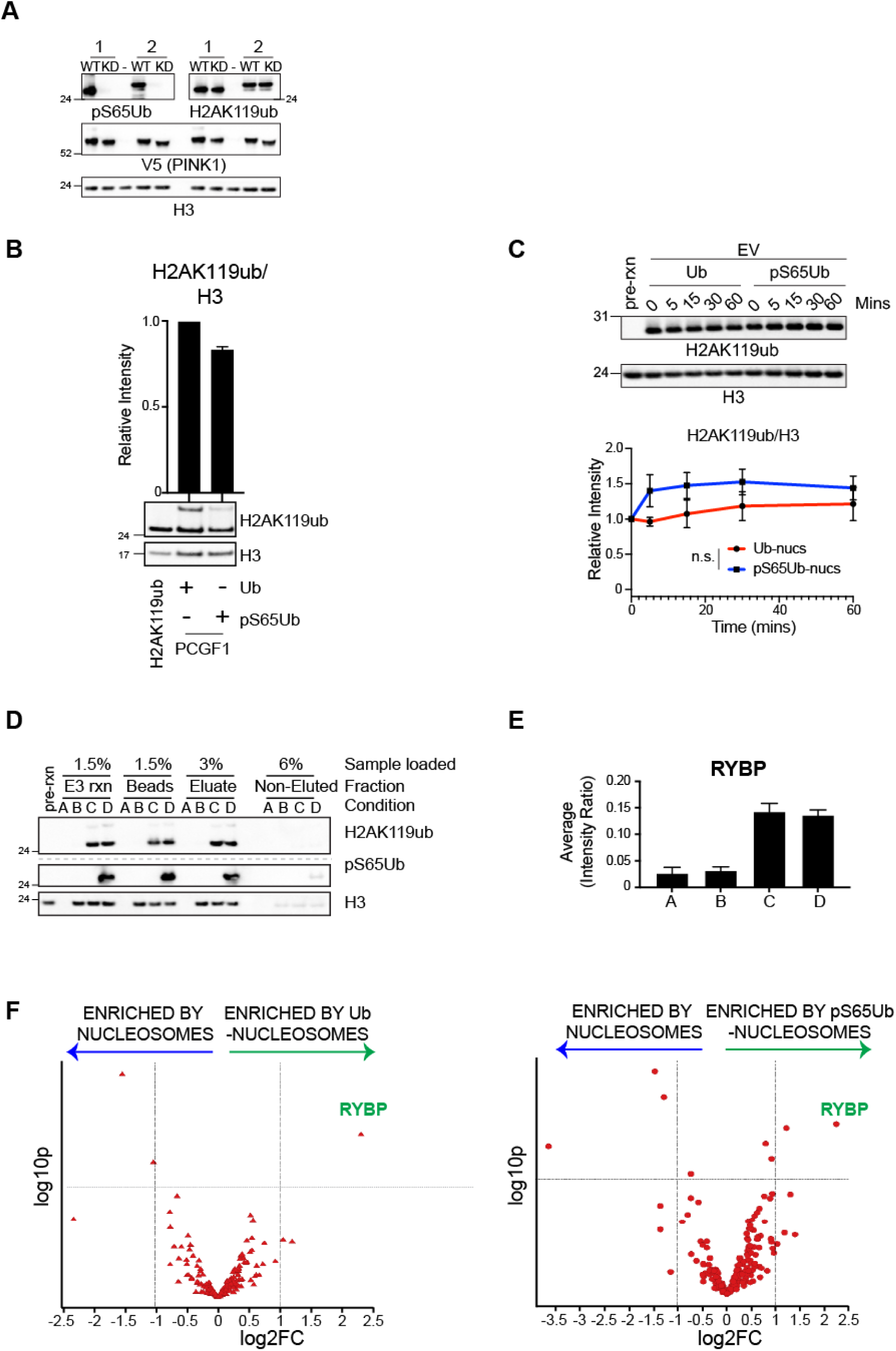
Characterizing pS65Ub-nucleosomes. **A** - Stably H2AK119-ubiquitinated nucleosomes (1) or nucleosomes *in vitro* ligated with FLAG-ubiquitin in-house (2) were subjected to *in vitro* kinase assays in the presence of WT or kinase dead (KD) insect Pink1-V5. Pre-ligation/kinase reaction nucleosome samples were loaded as a negative control (-). WT iPink1 enriches the pS65Ub histone band without affecting H2AK119ub antibody recognition. **B** - Nucleosomes were modified *in vitro* with ubiquitin or pS65Ub by using a six-fold lower RING1B-PCGF1 concentration than Figure 4C to slow the reaction rate. Reduced mono- and multimonoubiquitination reveals that pS65Ub inhibits E3 ligase activity. Quantification of H3 normalized H2AK119ub levels from two independent repeats is shown (mean +/-SEM). **C** - H2AK119-ubiquitinated (Ub) or -pS65 ubiquitinated (pS65Ub) mononucleosomes were incubated with negative control beads (HA resin incubated with EV-transfected cell lysates) for 0-60 mins as indicated. Mean +/- SEM relative H2AK119ub levels from four independent experiments are shown. The effect of ubiquitin type on reaction rate was not significant by two way ANOVA. **D** - Control samples for pS65Ub-nucleosome affinity-purification mass spectrometry experiment. Control mixes A (reaction mix without nucleosomes or E3) and B (reaction mix without E3), and both ubiquitinated (C) and pS65 ubiquitinated (D) mononucleosomes were prepared (E3 rxn), with pre-reaction nucleosomes loaded as a control for ubiquitination efficiency. Mixes were incubated with streptavidin beads to immobilise nucleosomes, then washed beads (Beads) were mixed with HeLa nuclear lysate to capture interacting proteins. After affinity purification and washing, bound proteins were eluted (Eluate), and prepped for TMT-based quantitative proteomics. Proteins retained on beads (Non-Eluted) were examined to confirm elution efficiency. Dashes lines indicate where identical samples were analysed on separate gels. **E** - TMT intensities for RYBP from samples A-D. Mean +/- SEM from three independent repeats. **F** - Volcano plot showing averaged protein intensities from unmodified vs ubiquitinated nucleosomes and unmodified vs pS65 ubiquitinated nucleosomes. Fold change and adjusted significance values are plotted logarithmically on the X and Y axis respectively. RYBP (annotated) is enriched in samples C and D. * <p0.05, ** <p0.01,*** <p0.001, **** <p0.0001, n.s. not significant.

**Extended Data Figure 6.**
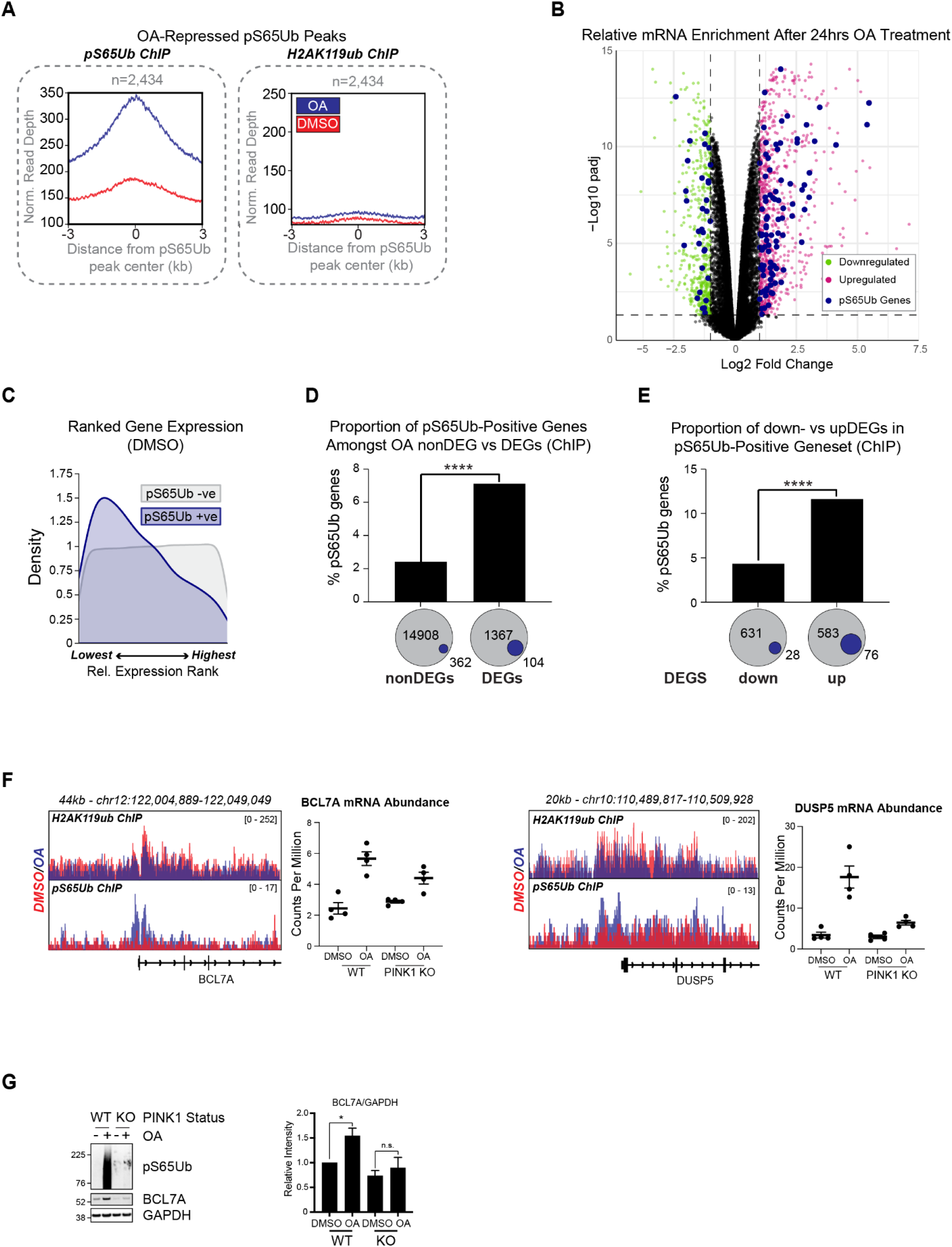
Evidence for pS65Ub-dependent transcriptional upregulation. **A** - Averaged pS65Ub and H2AK119ub ChIP Seq reads from DMSO (red) and OA (blue) conditions, centered on OA-repressed pS65Ub peaks, are depicted as metagene profiles. **B** - Volcano plot displaying RNA Seq results from WT HeLa cells treated with OA (1µM, 24 hrs, Extended Data Table 12). Genes significantly (adjusted p value <0.05) downregulated (log_2_(OA/DMSO)<-1, green) and upregulated (log_2_(OA/DMSO)>1, pink) after OA treatment are annotated. pS65Ub-positive genes (mean relative enrichment score >0.5, Extended Data Table 3) are overlaid in blue. **C** - RNA Seq results from HeLa cells treated with DMSO for 24 hrs displayed in a Kernel density estimate plot, with genes ranked by relative expression level and color coded based on presence (blue) or absence (grey) in the pS65Ub CUT&RUN geneset (mean relative enrichment score >0.5, Extended Data Table 3). **D** - pS65Ub-positive ChIP Seq genes (Extended Data Table 6) are overrepresented among OA-dependent DEGs in WT HeLa cells. Total pS65Ub-positive (blue) and negative (grey) DEG and non-DEG numbers listed below. Fisher’s Exact Test, odds ratio 2.48. **E** - The pS65Ub-positive ChIP Seq gene set contains significantly more OA-dependent upDEGs than downDEGs. Fisher’s Exact Test, odds ratio 2.94. **F** - Representative examples of candidate genes displaying both pS65Ub-dependent H2AK119ub depletion and gene expression. ChIP peak scale units = reads per kilobase per million. **G** - WT and PINK1 KO HeLa were treated with OA (1μM, 24 hrs) then whole-cell lysates were examined by immunoblot. Normalized BCL7A levels were quantified in three independent replicate experiments. Mean +/- SEM, * p<0.05, n.s. = not significant. **** <p0.0001.

**Extended Data Figure 7.**
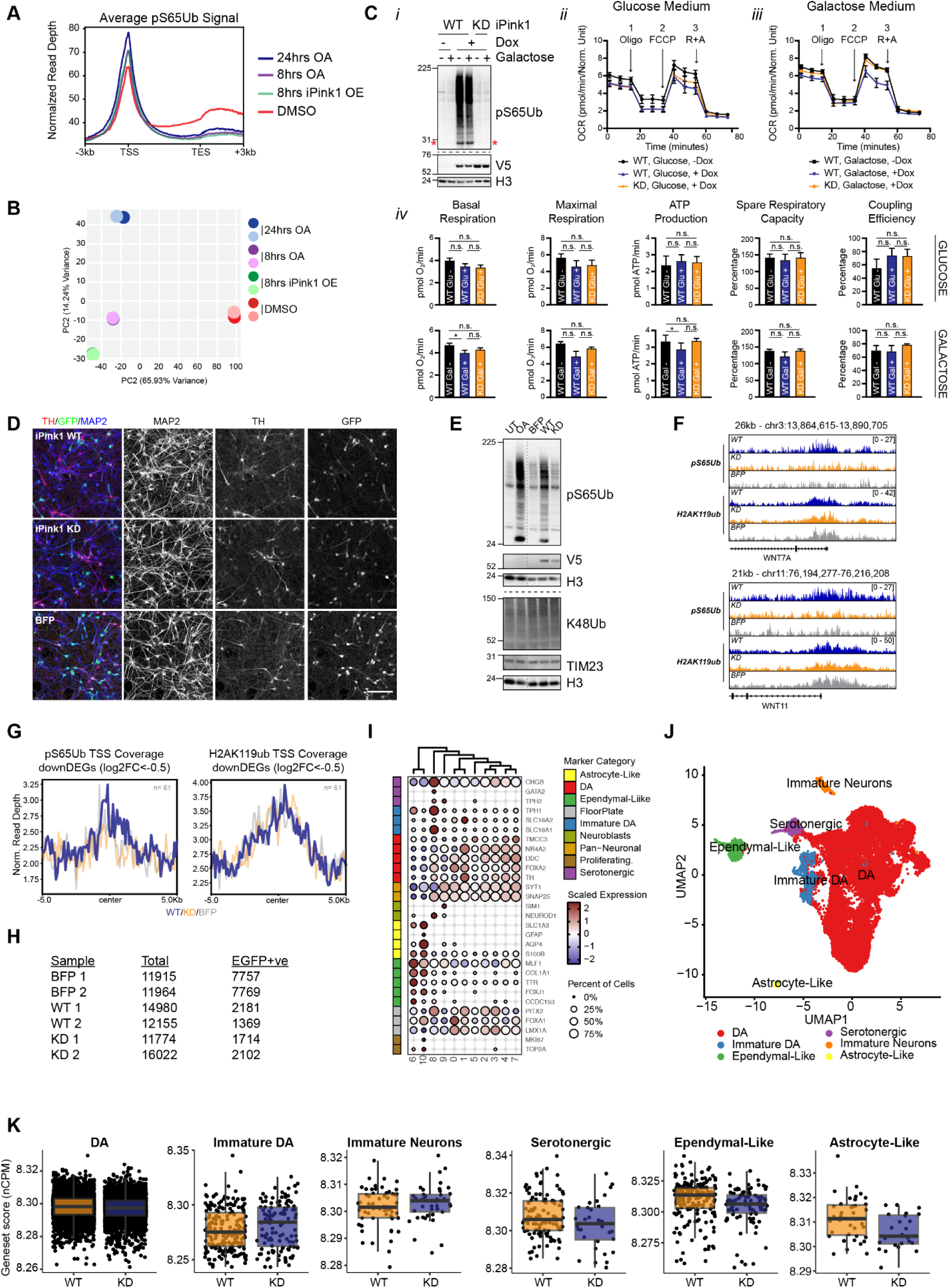
Characterizing pS65Ub-dependent phenotypes using the iPink1 model in dopaminergic neuron differentiation. **A** - Metagene profiles showing averaged pS65Ub ChIP Seq signal at the transcription start (TSS) and end sites (TES) of all annotated genes in HeLa cells treated with OA (8, 24 hrs), DMSO, or overexpressing WT iPink1 for 8 hrs. **B** - Principle component (PC) analysis plot comparing ChIP Seq signal in HeLa cells treated as indicated. **C** - HeLa cells dox-inducibly expressing WT or KD iPink1 were grown for 72 hours in media containing the standard glucose concentration used throughout this manuscript or in galactose media, which exacerbates mitochondrial dysfunction, before analysis. i) Immunoblotting shows pS65 ubiquitination only upon WT iPink1 expression. A red asterisk annotates the pS65Ub-histone band and a dashed line shows where identical samples were analysed on separate blots. ii/iii) Oxygen consumption rate (OCR - picomoles of oxygen per minute normalized to cell number) was measured over time in cells cultured as above following sequential addition of specific toxins at the indicated time points (1 = oligomycin, 2 = FCCP, 3 = rotenone + antimycin A). Mean +/- SEM from three independent replicates. iv) Quantitative analysis of key mitochondrial parameters derived from the OCR profiles. Mean +/- SEM from three independent repeats. **D** - Immunofluorescence images of dopaminergic (DA) neuron cultures transduced as precursors with iPink1 WT, iPink1 KD or BFP then stained for tyrosine hydroxylase (TH, red, DA marker), GFP (green, transduction marker) and MAP2 (blue, neuronal marker), scale bar = 50µm. **E** - iPSC-derived DA neurons were transduced as precursors with WT or KD iPink1, or BFP, or left untransduced. At maturity, untransduced cells were treated with OA (1µM, 24 hrs) or vehicle before Western blot analysis. **F** - Normalized pS65Ub and H2AK119ub ChIP signal in DA neuronal cultures expressing iPink1 WT/KD or BFP (scale units = reads per kilobase per million). Representative genes WNT7A and WNT11 were both significantly upregulated in bulk RNA Seq analysis (Extended Data Table 13) and are present in the MEISSNER_BRAIN_HCP_WITH_H3K4ME3_AND_H3K27ME3 MSigDB gene set. **G** - Metagene profiles showing averaged pS65Ub and H2AK119ub ChIP Seq signal at the transcription start site (TSS) of WT vs KD iPink1 downDEGs (Extended Data Table 13). **H** - Total and EGFP-positive cell numbers per condition as detected by scRNA Seq. **I** - Cell type marker gene expression in each of the major scRNAseq clusters, showing expression level and percentage of cells expressing each marker. **J** - UMAP plot of EGFP-positive cells from all conditions with predicted cell identities annotated. **K** - Box and whisker plot showing mean expression of genes in MSigDB gene set ‘MEISSNER_BRAIN_HCP_WITH_H3K4ME3_AND_H3K27ME3’ across each predicted major cell type. Units = normalized counts per million.

**Extended Data Figure 8.**
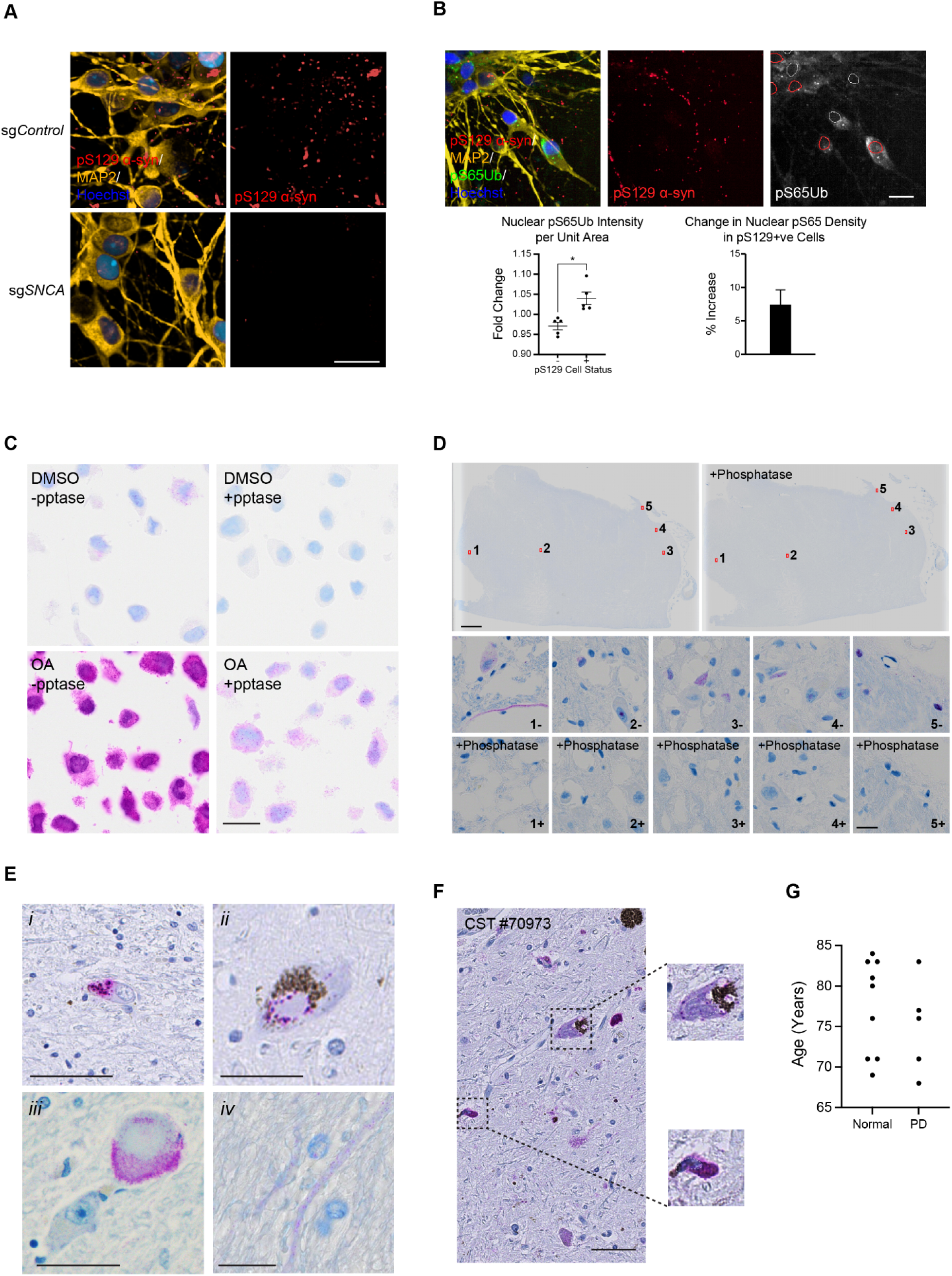
Validating pS65Ub immunohistological staining in human tissues. **A** - NGN2-cortical neurons expressing dCas9-KRAB were transduced with lentivirus encoding sgRNAs targeting SNCA or a non-targeting control (sgControl), then treated with PFFs for 14 days. Fixed cells were stained for pS129 α-Syn (red), MAP2 (yellow) and DNA (blue) and confocal images were taken. Representative images show the degree of pS129 α-Syn loss on α-synuclein knock down (sg*SNCA*) and demonstrate the effect of α-synuclein pre-formed fibril (PFF) seeding when α-synuclein expression is expressed to endogenous levels. **B** - NGN2-cortical neurons were treated with α-synuclein pre-formed fibrils (PFFs) for 14 days, fixed, then stained for pS129 α-synuclein (red), pS65Ub (green), MAP2 (yellow) and DNA (blue). In displayed confocal images, white (pS129-ve) and red (pS129+ve) outlines denote nuclear boundaries, contracted inward to yield conservative boundary estimates. Quantifications shows nuclear pS65Ub density in cells with and without pathological α-synuclein aggregation (+/- pS129 α-syn), expressed as a proportion of the total cell population (left) and as the percentage increase in pS129+ve vs -ve cells (right). Five independent differentiation repeats were quantified, mean +/- SEM. **C** - HeLa treated with OA (1µM,18 hrs) were fixed, sectioned, subject to lambda phosphatase treatment, then stained for pS65Ub (pink) and DNA (blue). Representative images show phosphatase-dependent clearance of pS65Ub signal. Scale bar 20µm. **D** - PD SNpc sections from the same tissue block were treated +/- lambda phosphatase then stained as above. Top - full tissue samples with numbered regions showing position of each magnified inset, scale bar 200µm. Bottom - Magnified insets showing representative pS65Ub signal distribution, scale bar 20µm. **E** - Representative images of major non-nuclear pS65Ub distributions observed in PD SNpc: i) glia - predicted mitochondrial; ii) neuron - predicted mitochondrial; iii) neuron: predicted Lewy body; iv) neuron: axonal. Scale bars 20µm. **F** - Representative image of PD SNpc showing similar pS65Ub distributions with an independent antibody (Cell Signaling Technologies #70973). Black dashed boxes indicate regions magnified in inset, dashed circles outline proposed nuclear area. Scale bar 50µm. **G** - Distributions of age at collection of healthy and PD post-mortem samples used in Figure 7B/C.

**Extended Data Figure 9.**
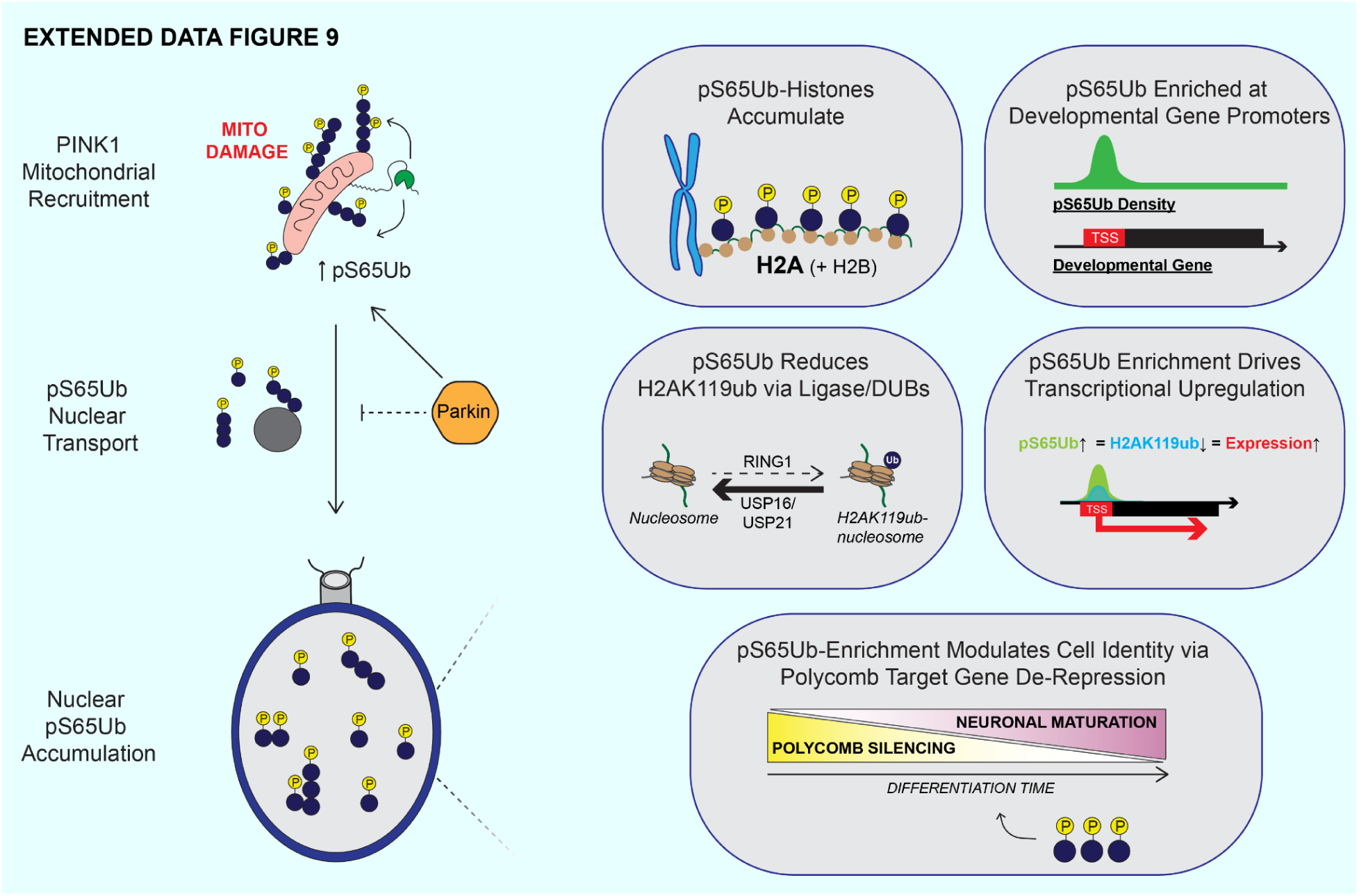
pS65Ub as a Secondary Messenger -. The proposed mechanisms for pS65Ub nuclear accumulation and H2AK119-dependent transcriptional regulation are depicted schematically.

**Extended Data Table 1 - Nuclear pS65Ub affinity purification mass spectrometry results.** Relative TMT enrichment scores for proteins identified in nuclear pS65Ub pulldown experiment (Figure 1C, D). Four comparisons were made, listed in the comparison column: wild type OA vs vehicle/DMSO (WTOA_vs_WTVeh), wild type OA vs PINK1 KO OA (WTOA_vs_KOOA), PINK1 knockout OA vs DMSO (KOOA_vs_KOVeh) and wild type DMSO vs PINK1 KO DMSO (WTVeh_KOVeh). Protein, log_2_(fold change), standard error, p and adjusted p values for all comparisons are shown. Candidate pS65Ub-modified proteins are marked in the final column with an asterisk.

**Extended Data Table 2 - Nuclear pS65Ub affinity purification mass spectrometry gene set enrichment results.** Gene Set enrichment analysis (GSEA) on candidate nuclear pS65Ub targets identified Extended Data Table 1. Annotated pathway, normalized enrichment score (NES), p and adjusted p values are shown. GSEA pathways significantly enriched (adjp<0.05) in WTOA_vs_WTVeh or WTOA_vs_KOOA comparisons are marked with an asterisk in the final column.

**Extended Data Table 3 - pS65Ub CUT&RUN geneset.** Genes displaying OA-dependent pS65Ub enrichment in the region spanning +/- 1kb of the transcription start site are shown. Relative enrichment (RE) scores (ranked peak score equivalent to -log10(p value) from MACS2 peak calling) from three independent repeats are shown along with mean relative enrichment score.

**Extended Data Table 4 - Gene set enrichment analysis results from pS65Ub CUT&RUN geneset.** GSEA was performed on genes showing pS65Ub signal enrichment at promoters after 24 hrs OA treatment (Extended Data Table 3). Term name, p and adjusted p values, the % of genes matching each term and the gene identities are listed.

**Extended Data Table 5 - pS65Ub coregulator analysis results.** The gene list of TSS-proximal pS65Ub CUT&RUN peaks from Extended Data Table 3 was used as input to the Binding Analysis for Regulation of Transcription (BART) algorithm^66^, which infers corregulators of input gene lists. Max_auc refers to the maximum association score in multiple chromatin profiling datasets for each protein. Results were derived using default parameters from the web server: http://bartweb.org/.

**Extended Data Table 6 - Relative pS65Ub ChIP Seq peak enrichment at TSS region in HeLa cells treated with 24 hrs OA vs DMSO.** Genes displaying significant pS65Ub ChIP Seq signal enrichment at the TSS region in HeLa cells after 24 hrs OA treatment are shown. Log_2_ fold change, log_2_ fold change standard error, p value and adjusted p value are given. An asterisk in the final column indicates where the same genes were significantly enriched in neurons (Extended Data Table 7).

**Extended Data Table 7 - Relative pS65Ub ChIP Seq peak enrichment at TSS region in Neurons treated with 24 hrs OA vs DMSO.** Genes displaying significant pS65Ub ChIP Seq signal enrichment at the transcription start site (TSS) region in neurons after 24 hrs OA treatment are listed. Log_2_ fold change, log_2_ fold change standard error, p value and adjusted p value are given. An asterisk in the final column indicates where the same genes were significantly enriched in HeLa cells (Extended Data Table 6).

**Extended Data Table 8 - Gene set enrichment analysis results from pS65Ub ChIP gene set (HeLa).** GSEA was performed on pS65Ub-enriched genes identified in OA treated HeLa cells (Extended Data Table 6).

**Extended Data Table 9 - Gene set enrichment analysis results from pS65Ub ChIP gene set (Neurons).** GSEA was performed on pS65Ub-enriched genes identified in OA treated neurons (Extended Data Table 7).

**Extended Data Table 10 - Gene set enrichment analysis results from pS65Ub-positive genes enriched in both neurons and HeLa cells.** GSEA was performed on pS65Ub targets shared between neurons and HeLa cells (Extended Data Tables 7, 8).

**Extended Data Table 11 - Nuclear pS65Ub affinity purification mass spectrometry results.** Relative TMT enrichment values for proteins identified in pS65Ub-nucleosome APMS experiment. Four comparisons were made, listed in the comparison column: phosphoubiquitinated vs ubiquitinated nucleosomes (NucS65Ub_NucWTUb), phosphoubiquitinated vs non-ubiquitinaed nucleosomes (NucS65Ub_Nuc), ubiquitinated vs non-ubiquitinated nucleosomes (NucWTUb_Nuc) and non-ubiquitinated nucleosomes vs beads alone (Nuc_Beads). Proteins are listed in each row, with log_2_ fold change, standard error, and both p and adjusted p values for all comparisons shown.

**Extended Data Table 12 - Differential gene expression analysis of HeLa cells treated with oligomycin + antimycin.** Wild type and PINK1 knockout HeLa cells were treated with OA for 24 hrs before RNA Seq analysis. Each row represents a gene identified in the analysis, with OA versus DMSO log_2_ fold change value, p value and adjusted p value from four independent repeats in each cell type (WT and KO). PINK1-dependent relative OA-induced expression was calculated by subtracting log_2_(OA/DMSO) fold change values in KO from WT conditions (dFC). Genes identified in the pS65Ub CUT&RUN experiment (mean relative enrichment score >0.5, Extended Data Table 3) are identified with an asterisk.

**Extended Data Table 13 - Differential gene expression analysis of dopaminergic neurons transduced with wild type or kinase dead iPink1.** Dopaminergic neurons precursors were transduced with lentiviruses expressing wild type (WT) or kinase dead (KD) iPink1 on differentiation day 22 before bulk RNA Seq analysis on day 62. Each row represents a gene identified in the analysis, with WT versus KD log_2_ fold change value, p value and adjusted p value from 3 independent repeats.

**Extended Data Table 14 - Differential gene expression analysis of dopaminergic neurons transduced with wild type iPink1 or BFP.** Dopaminergic neurons precursors were transduced with lentiviruses expressing BFP or wild type (WT) iPink1 on differentiation day 22 before bulk RNA Seq analysis on day 62. Each row represents a gene identified in the analysis, with WT versus BFP log_2_ fold change value, p value and adjusted p value from 3 independent repeats.

**Extended Data Table 15 - Differential gene expression analysis of dopaminergic neurons transduced with kinase dead iPink1 or BFP.** Dopaminergic neurons precursors were transduced with lentiviruses expressing BFP or kinase dead (KD) iPink1 on differentiation day 22 before bulk RNA Seq analysis on day 62. Each row represents a gene identified in the analysis, with KD versus BFP log_2_ fold change value, p value and adjusted p value from 3 independent repeats.

